# Conduction velocity along the local axons of parvalbumin interneurons correlates with the degree of axonal myelination

**DOI:** 10.1101/2020.10.10.334656

**Authors:** Kristina D. Micheva, Marianna Kiraly, Marc M. Perez, Daniel V. Madison

## Abstract

Parvalbumin-containing (PV+) basket cells in mammalian neocortex are fast-spiking interneurons that regulate the activity of local neuronal circuits in multiple ways. Even though PV+ basket cells are locally projecting interneurons, their axons are myelinated. Can this myelination contribute in any significant way to the speed of action potential propagation along such short axons? We used dual whole cell recordings of synaptically connected PV+ interneurons and their postsynaptic target in acutely-prepared neocortical slices from adult mice to measure the amplitude and latency of single presynaptic action potential-evoked inhibitory postsynaptic currents (IPSCs). These same neurons were then imaged with immunofluorescent array tomography, the synaptic contacts between them identified and a precise map of the connections was generated, with the exact axonal length and extent of myelin coverage. Our results support that myelination of PV+ basket cells significantly increases conduction velocity, and does so to a degree that can be physiologically relevant.

---

Parvalbumin-containing basket cells comprise fewer than 8% of neocortical neurons (Rudy et al. 2011), yet they control a multitude of neuronal circuit functions with high reliability and exquisite temporal precision. PV+ basket cells are fast-spiking interneurons (Hu et al. 2014), and each one of them contacts hundreds of pyramidal neurons in its vicinity (Packer et al. 2011), with synaptic connections strategically positioned for strong influence on the cell bodies and proximal dendrites (Somogyi et al. 1983; Tamás et al. 1997; Kawaguchi and Kubota 1998). Their electrophysiological behavior is supported by numerous molecular and structural specializations (Hu et 1. 2014), including high Na+ channel density in their axons (Hu and Jonas 2014), fast presynaptic Ca2+ channels of the P/Q type (Zaitsev et al. 2007), and high mitochondrial content (Gulyás et al. 2006; Cserép et al. 2018). Even before it was known that these interneurons express PV (Celio 1986; Kawaguchi et al. 1987), classical EM studies that identified these basket cells by morphological and electrophysiological criteria (Somogyi et al. 1983; DeFelipe et al. 1986, Somogyi and Soltész 1986) showed that their axons are myelinated and hypothesized that this would result in higher conduction velocity and reliability of neurotransmission. However, more recent studies revealing that myelin consistently covers only the proximal axonal arbor and in a patchy manner (Micheva et al, 2016; Stedehouder et al. 2017), cast doubt on whether myelin in this case is indeed speeding up action potential propagation along the rather short distances to their local targets, suggesting other roles for myelination, such as energy conservation, trophic support or potential for plasticity.

Here we combined electrophysiological measurements with array tomographic mapping of PV+ basket cells connections to directly measure the time that elapses between the firing of a presynaptic action potential in a PV+ interneuron, and the onset of the inhibitory postsynaptic current it triggers, and to correlate this with the measured axonal length including the fraction of that axon that is myelinated. Basket cells from a variety of ages and cortical layers of the adult mouse medial temporal lobe (Beaudin et al. 2013) were analyzed aiming to capture a broad range of myelin coverage. We performed simultaneous whole cell recordings between identified PV+ interneurons and monosynaptically-efferent to postsynaptic target neurons in mouse neocortex. During the electrophysiological recordings, the two neurons, presynaptic interneuron and postsynaptic target neurons (either a pyramidal neurons or another interneuron) were dye-filled through the recording electrode. Immediately after the termination of the physiological recordings, slices were fixed and processed for array tomography (AT), which was used to analyze the synaptic connections and axonal paths, including myelination, using the injected dye and immunofluorescent markers to identify the recorded neurons and their processes. AT is based on digital reconstruction of images acquired from arrays of serial ultrathin sections (100 nm in the present study) which allows light level identification and tracing of individual axons and their myelin sheath (Micheva et al. 2016), as well as synapses (Micheva et al. 2010; Collman et al. 2014). Knowing the precise length of the axonal paths that connect two neurons in a pair, as well as the latency of the postsynaptic response allowed us to calculate the conduction velocity and explore its relation to myelin. We found that conduction velocity significantly correlated with the extent of axon myelination, consistent with the role of myelin in speeding up neurotransmission of PV+ basket cells. In addition, we observed a unique mesh-like connectivity pattern between individual PV+ basket cells and their postsynaptic partner, with multiple axonal paths contacting the postsynaptic neuron at multiple subcellular domains. In particular, the targeting of different axonal paths to the same postsynaptic domain was a salient feature in pairs with high conduction velocity, suggesting that this mesh-like connectivity pattern is an additional specialization of PV+ basket cells supporting their functions in neuronal circuits.

## Materials and Methods

### Animals

Thirty-one C57BL/6 mice, ages 1 to 7 months, were used for this study. All procedures related to the care and treatment of animals were carried out in accordance with the Guide for the Care and Use of Laboratory Animals of the National Institutes of Health and approved by the Administrative Panel on Laboratory Animal Care at Stanford University.

### Preparation of Acute Neocortical Slices

Brain slices were prepared exactly following the protocol of Ting et al., 2018. Our attempted variations on this protocol tended to decrease the quality and viability of the slices. Briefly, mouse brains were hemisected, placed into warm 2% agarose, which was then hardened by ice-chilling and inserted into the feed tube of a Compresstome vibrating microtome (Precisionary Instruments), and cut to a thickness of 300 μm. Slices were stored submerged for at least one hour at room temperature in a Model 4 Brain Slice Keeper (AutoMate Scientific). During this incubation period, slices were subjected to the Wisteria floribunda lectin staining procedure below. For recording, slices were transferred to a submerged/superfusing slice chamber with a glass coverslip bottom (Warner Instruments) on the stage of an upright microscope.

### PV+ Interneuron identification

To identify PV+ interneurons in living tissue, the perineuronal nets that selectively surround these neurons were stained in the live slices using fluorescein labeled Wisteria floribunda lectin (Vector Labs FL-1351), following the protocol of Hoppenrath et al., 2016. Briefly, immediately after sectioning on the Compresstome, slices were transferred into a small volume of holding buffer (in mM: 92 NaCl, 2.5 KC1, 1.25 NaH_2_PO_4_, 30 NaHCO_3_, 20 HEPES, 25 glucose, 2 thiourea, 5 Na-ascorbate, 3 Na-pyruvate, 2 CaCl_2_H_2_O, and 2 MgSO_4_·7H_2_O, pH to 7.3–7.4) containing fluorescein labeled Wisteria floribunda lectin (20 μg/ml) and maintained at room temperature under a 95% O_2_, 5 % CO_2_ atmosphere for 1 hour, before either being transferred directly into the recording chamber, or kept in the slice keeper for recording later in the day. Labeled perineuronal nets were detected in the live slices using epiflourescence (Nikon GFP filter cube).

In preliminary experiments, the identity of the neurons labeled by the Wisteria floribunda lectin was confirmed by subsequent immunofluorescent labeling with anti-Parvalbumin antibody (Figure 1A). For these preliminary experiments, the slices were chemically fixed with 2% formaldehyde/2% glutaraldehyde solution in PBS as outlined below, then washed in PBS, and permeabilized for 2 h in PBS containing 0.04% Triton-X and 0.02 % DMSO. They were then incubated in blocking solution (1% BSA) overnight at room temperature, then in primary antibody (SWANT PV27, 1:1000) for 10 days at 4°C, washed in the permeabilizing solution for 1 day, and incubated in the secondary antibody (goat anti-rabbit Alexa Fluor 488, 1:200) and Streptavidin-Alexa 594 (ThermoFisher Scientific S11227, 1:100) for 6 days at 4°C.

**Figure 1.**
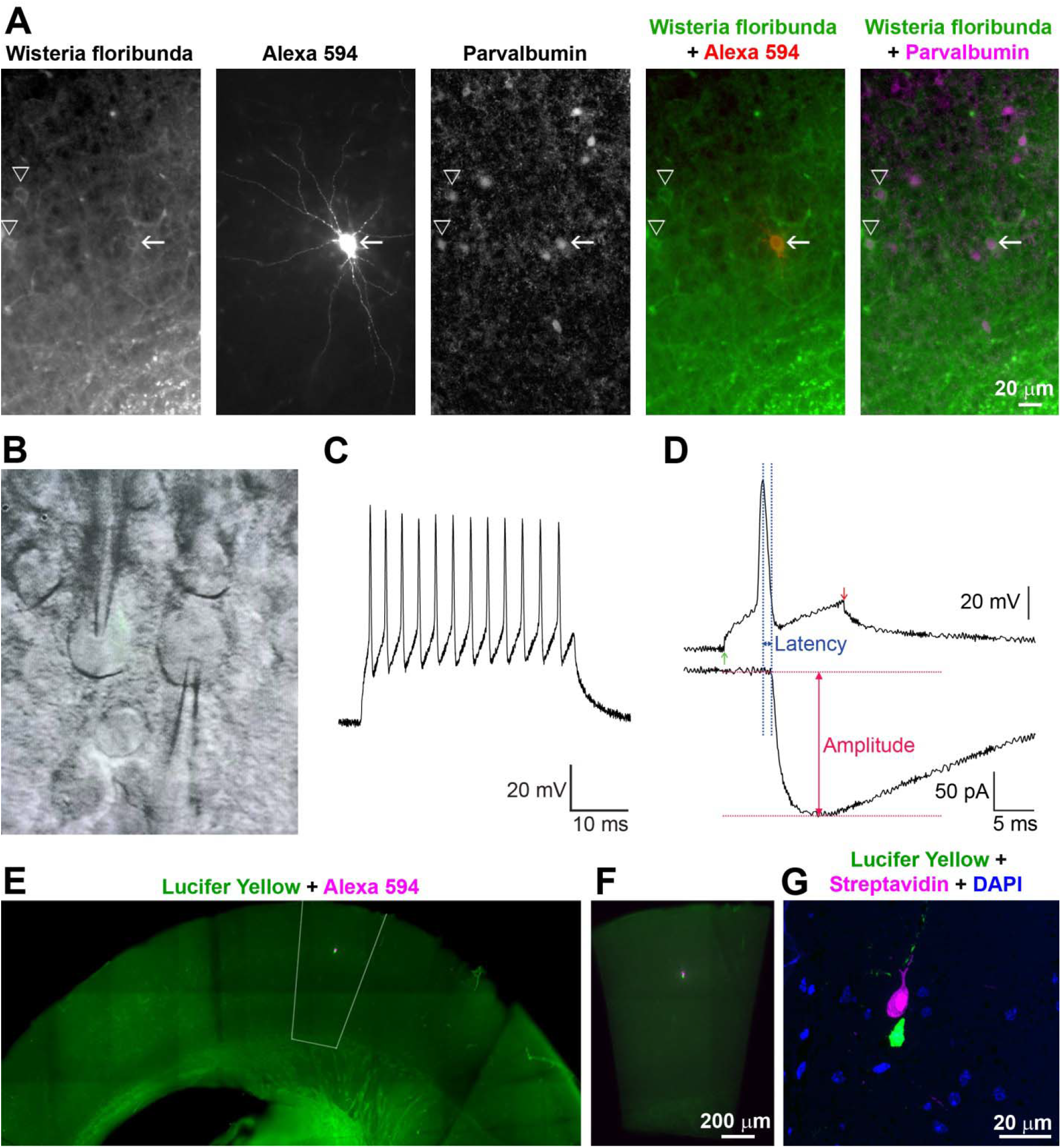
Experimental approach. **A.** Preliminary experiments confirming the identification of PV+ interneurons in mouse acute cortical slices. Wisteria floribunda lectin selectively labels perineuronal nets in live slices. The neuron filled with Alexa 594 hydrazide (arrow) is PV immunopositive, as are the other neurons surrounded by labeled perineuronal nets (arrowheads). Because Wisteria floribunda labels only the surface layer of the slice (approximately 50 μm depth), there are other PV+ neurons without stained perineuronal nets. **B.** Differential interference contrast image of a paired recording from a mouse acute neocortical slice with both electrodes visible (photographed off a video monitor used to locate cells during the experiment). **C.** Action potential repetitive firing pattern of a PV+ interneuron in response to a prolonged current injection, and the resultant synaptic currents in the postsynaptic neuron. **D.** Analysis of the amplitude and latency of the postsynaptic response. To aid in consistent measurement of latency, the peak of the presynaptic action potential was used as the starting time point. **E.** Fluorescence image from an acutely-prepared mouse neocortical slice after chemical fixation, showing the position of the neuronal pair with the presynaptic neuron filled with Lucifer Yellow and the postsynaptic neuron filled with Neurobiotin and Alexa Fluor 594 hydrazide. The outlined area was dissected out for further processing. **F.** The dissected out area is centered on the recorded pair and encompasses all cortical layers. **G.** An ultrathin section (100 nm) through the same neuronal pair as in E and F, imaged after embedding and ultrathin sectioning. The presynaptic neuron is visualized with Streptavidin 594 detecting the Neurobiotin fill, and the postsynaptic neuron – with Lucifer Yellow immunolabeling.

### Electrophysiology

We performed simultaneous whole-cell recordings from two different neurons, a presynaptic cell being an indentified PV+ basket cells, and a postsynaptic cell being a nearby unlabeled neuron. In most cases, the postsynaptic neuron was a pyramidal neuron, but sometimes, it was an interneuron. The presynaptic interneuron was recorded in current clamp, and the postsynaptic neuron in voltage clamp. Single action potentials were elicited from the presynaptic interneuron by the injection of positive current through the recording electrode, amplitude of the current pulse adjusted to produce a single action potential. The postsynaptic recording was examined for time-locked postsynaptic responses to that presynaptic action potential. We measured the amplitude of the evoked postsynaptic current and the latency of that current, relative in time to the peak of the presynaptic action potential (Figure 1). The detailed methods, including recording configurations, solutions, electrodes, etc. were as in (Pavlidis & Madison, 1999, Montgomery et al., 2001), except that the postsynaptic electrode internal solution consisted of (in mM): 120 KCl, 40 HEPES, 2 Mg-ATP, 0.317 Na-GTP, and 10.7 MgCl_2_. Besides making the IPSCs larger and easier to detect, use of high chloride solution also reversed the usual polarity of inhibitory synaptic currents to inward. To mark the neurons for subsequent AT analysis, the internal electrode solution included 0.1% Alexa Fluor 594 hydrazide (ThermoFisher Scientific A-10438) and 0.5% Neurobiotin (Vector Laboratories SP-1120-50) for the presynaptic PV+ interneuron, and 0.2% Lucifer Yellow CH, potassium salt (ThermoFisher Scientific L1177) for the postsynaptic neuron. At the end of each electrophysiological experiment, the cortical slice was removed from the recording chamber and chemically fixed.

### Slice fixation

At the end of the experiment, the slice was placed in a fixative consisting of 2% formaldehyde (diluted from 8% formaldehyde, EM grade, Electron Microscopy Sciences 157-8) and 2 % glutaraldehyde (diluted from 25% glutaraldehyde, EM grade, Electron Microscopy Sciences 16220) in PBS, for 1 hour at room temperature, followed by 24h at 4°C.

### Embedding in resin

After chemical fixation, slices were washed in PBS, and a smaller area (approximately 1 by 2 mm, Figure 1E, F) around the labeled neurons was dissected out and dehydrated serially in washes of 50%, 70%, 70% ethanol. Each step was done for 10 min at 4°C. The dehydrated tissue was then infiltrated with LR White resin, hard grade (Electron Microscopy Sciences 14383), first with a mixture of 70% ethanol and LR White (1:3) and then in three changes of 100% LR White, 10 min at room temperature each step. The slice was left in unpolymerized LR White resin overnight at 4°C, then transferred to a gelatin capsule size “00” (Electron Microscopy Sciences 70100) and polymerized for 24h at 55°C. The polymerized blocks with tissue were stored in the dark at room temperature.

### Sectioning

The blocks, once removed from the gelatin capsule, were trimmed around the tissue to the shape of a trapezoid approximately 2 mm wide and 0.5 mm high, and glue (Weldwood Contact Cement diluted with xylene) was applied with a thin paint brush to the leading and trailing edges of the block pyramid. The embedded plastic block was cut on an ultramicrotome (Leica Ultracut EM UC6) into 100-nm-thick serial sections, which were mounted on gelatin-coated coverslips. Typically, 1,000 to 1,200 serial sections containing the fixed tissue slice were obtained from each block.

### Immunofluorescent array tomography

Coverslips with sections containing the filled neurons were identified by visual inspection using a 10x objective under the fluorescence microscope, and were processed for standard indirect immunofluorescence, as described in Micheva et al. 2010. Antibodies were obtained from commercial sources and are listed in Table 1. The sections were pretreated with sodium borohydride [1% in Tris-buffered saline (TBS), pH 7.6 for 3 min] to reduce non-specific staining and autofluorescence. After a 20 min wash with TBS, the sections were incubated in 50 mM glycine in TBS for 5 min, followed by blocking solution (0.05% Tween 20 and 0.1% BSA in TBS) for 5 min. The primary antibodies were diluted in blocking solution as specified in Table 1 and were applied overnight at 4°C. After a 15-min wash in TBS, the sections were incubated with Alexa Fluor dye-conjugated secondary antibodies, highly cross-adsorbed (ThermoFisher Scientific), diluted 1:150 in blocking solution for 30 min at room temperature. Neurobiotin was detected using Streptavidin-Alexa 594 (ThermoFisher Scientific S11227), diluted at 1:200, and applied together with the secondary antibodies. Finally, sections were washed with TBS for 15 min, rinsed with distilled water, and mounted on glass slides using SlowFade Diamond Antifade Mountant with DAPI (ThermoFisher Scientific S36964). After the sections were imaged, the antibodies were eluted using a solution of 0.2 M NaOH and 0.02% SDS, and new antibodies were reapplied on some of the coverslips, as needed. The first round of staining always included anti-LY and Streptavidin-Alexa 594 to visualize the filled neurons and anti-MBP to detect myelin.

**Table 1.**
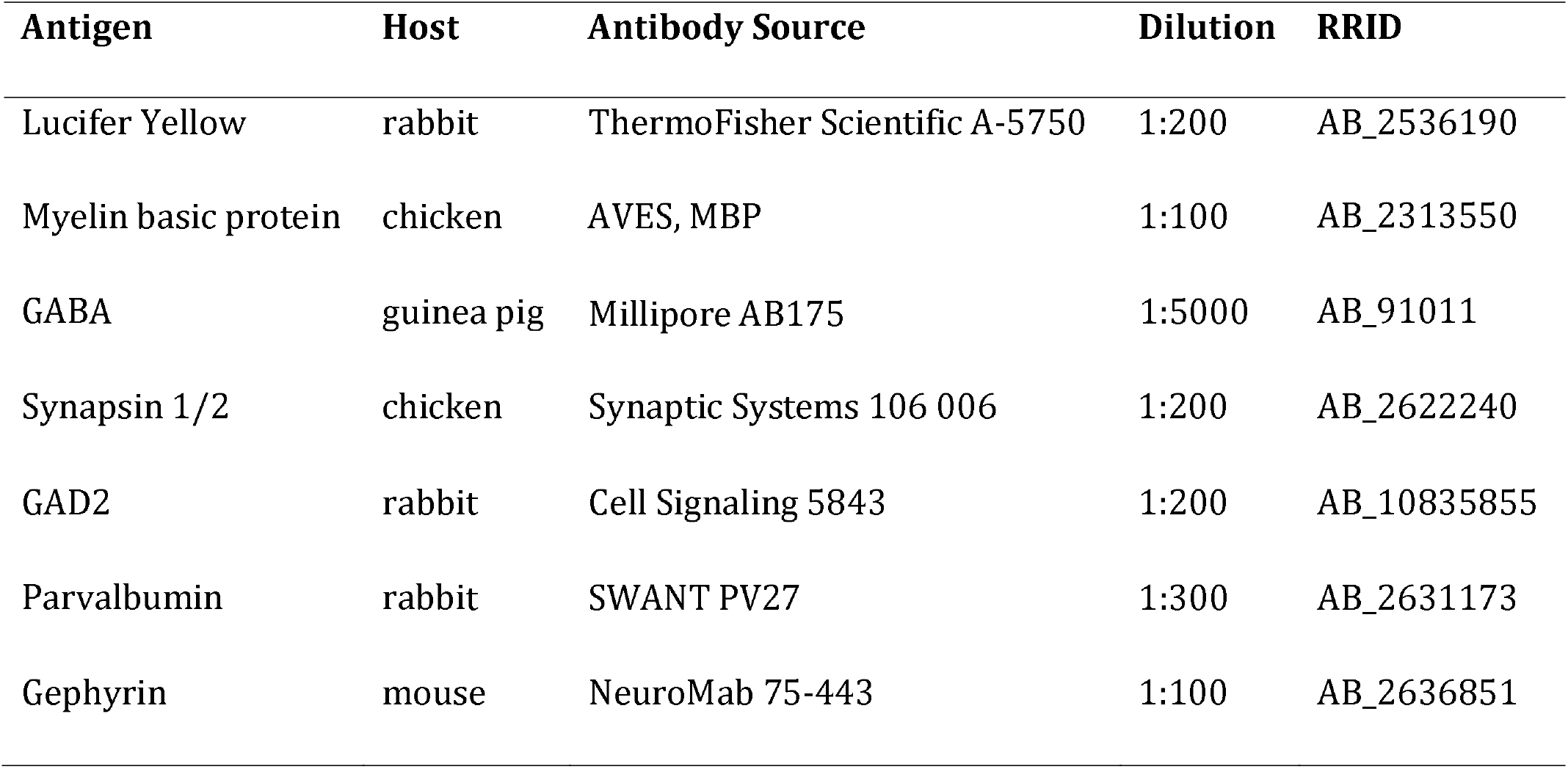
Antibodies used in the study.

| Antigen | Host | Antibody Source | Dilution | RRID |
| --- | --- | --- | --- | --- |
| Lucifer Yellow | rabbit | ThermoFisher Scientific A-5750 | 1:200 | AB_2536190 |
| Myelin basic protein | chicken | AVES, MBP | 1:100 | AB_2313550 |
| GABA | guinea pig | Millipore AB175 | 1:5000 | AB_91011 |
| Synapsin 1/2 | chicken | Synaptic Systems 106 006 | 1:200 | AB_2622240 |
| GAD2 | rabbit | Cell Signaling 5843 | 1:200 | AB_10835855 |
| Parvalbumin | rabbit | SWANT PV27 | 1:300 | AB_2631173 |
| Gephyrin | mouse | NeuroMab 75-443 | 1:100 | AB_2636851 |

### Imaging

The immunostained ribbons of sections were imaged on an automated epifluorescence microscope (Zeiss AxioImager Z1) using a 63x Plan-Apochromat 1.4 NA oil objective. To define the position list for the automated imaging, a custom Python-based graphical user interface, MosaicPlanner (obtained from https://code.google.com/archive/p/smithlabsoftware/), was used to automatically find corresponding locations across the serial sections. On average, 515 sections per neuron pair were imaged (ranging between 264 to 783 sections), resulting in volumes of approximately 140 μm by 400 μm by 51.5 μm centered on the filled neurons. Images from different imaging sessions were registered using a DAPI stain present in the mounting medium. The images from the serial sections were also aligned using the DAPI signal. Both image registration and alignment were performed with the MultiStackReg plugin in FIJI (Schindelin et al. 2012).

### Contact detection and axon tracing

A contact between the two filled neurons was identified as a direct apposition between an enlargement in the presynaptic axon and the postsynaptic neuron, which was present on two or more consecutive sections. Thirty-five contacts were subsequently immunostained with synaptic markers to verify whether contacts selected by these criteria were indeed synapses. Using Fiji, the axons were traced back from the synaptic contacts to the neuronal cell body. The myelinated internodes were identified using the MBP immunofluorescence. The following measurements were obtained: total axonal distance to each contact, length and diameter of axon initial segment, length, diameter and number of myelinated internodes, length of nodes of Ranvier, distance of the postsynaptic target from the postsynaptic cell body. Because tissue dehydration and embedding with our protocol result in approximately 23% linear shrinkage, these measurements were adjusted accordingly (Busse and Smith 2013).

### Estimation of release probability at single synapses

Knowing both the rate at which all ‘n’ synapses in a given connection fail in the same trial, and the number of synapses comprising that connection, allowed us to calculate an estimate of the failure rate of single synapses, and by extension the release probability of single inhibitory synapses. Assuming that the probability is the same for all events, and that the events are independent:

With n = number of synapses in the connection, y = the measured rate at which all n synapses fail in the same trial, and x being the unknown single synapse failure rate,

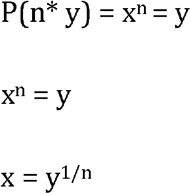

### Statistics

Measurements are presented as average ± standard deviation. For comparison between groups we used two-tailed unpaired t-test. Linear correlations between 2 variables were done using Pearson’s correlation coefficient. To prepare the boxplots, the web application BoxPlotR was used (http://boxplot.tyerslab.com; Spitzer et al. 2014).

## Results

Thirty-one synaptically-connected pairs of neurons were recorded in freshly-prepared (“acute”) neocortical slices taken from mouse medial temporal lobe (perirhinal and entorhinal cortex). In all cases, the presynaptic neuron was a PV+ interneuron. Fourteen of these pairs, where both neurons were brightly labeled such that their distal dendrites and axons could be followed, were further processed for AT analysis. In 11 out of the 14 pairs, the postsynaptic neuron was identified by its morphology as neocortical pyramidal neuron, the others being interneurons (Table 2). The electrophysiological recordings of pairs that were not adequately filled for AT analysis were included in the electrophysiological analysis only.

**Table 2.** Neuronal pairs analyzed by AT.

| Pair | Age (mo) | # sections | Cortical region | Layer | Postsyn. cell | IPSC (pA) | Latency (ms) | Failure rate % |
| --- | --- | --- | --- | --- | --- | --- | --- | --- |
| MK190129-1 | 1 | 413 | Perirhinal | 3/4 | Pyramidal | 344.16 | 1.06 | 0 |
| MK181212 | 1 | 727 | Perirhinal | 5 | Pyramidal | 34.966 | 1.23 | 14.29 |
| MK181213 | 1 | 587 | Entorhinal | 3/4 | Pyramidal | 14.33 | 1.01 | 6.25 |
| MK181217 | 1 | 548 | Entorhinal | 5 | Pyramidal | 35.73 | 1.35 | 12 |
| MK181211 | 1 | 783 | Perirhinal | 5/6 | Pyramidal | 34.73 | 1.02 | 0 |
| MK180814 | 2 | 420 | Perirhinal | 5/6 | Interneuron | 274.66 | 1.75 | 0 |
| MK190115-1 | 2 | 289 | Perirhinal | 3/4 | Pyramidal | 13.05 | 2.63 | 16 |
| MK190110 | 2 | 394 | Perirhinal | 4/5 | Pyramidal | 33.8 | 1.76 | 12.5 |
| MK190227 | 2 | 661 | Perirhinal | 4 | Pyramidal | 300.08 | 1.38 | 0 |
| MK180817-1* | 2 | 264 | Entorhinal | 5 | Pyramidal | 223.44 | 2.46 | 0 |
| MK181220 | 7 | 463 | Perirhinal | 3/4 | Interneuron | 377.39 | 0.93 | 0 |
| MK181127 | 7 | 424 | Perirhinal | 5 | Interneuron | 61.8 | 0.87 | 0 |
| MK181221-1 | 7 | 485 | Perirhinal | 4 | Pyramidal | 417.06 | 1.32 | 0 |
| MK190109-2 | 7 | 748 | Entorhinal | 5 | Pyramidal | 21.06 | 1.64 | 48.48 |

\* In this case, because the 2 neurons in the pair were much further apart along the z-axis than in the other pairs, two separate volumes were imaged – one centered on the presynaptic neuron (64 sections) and another one on the postsynaptic neuron (200 sections). Axonal paths could not be traced for this pair.

### Electrophysiological characterization of the interneuron connections

The average amplitude of the presynaptic action potential-evoked inhibitory postsynaptic current (IPSC) across all trials in each pair was calculated (excluding trials with synaptic failures in those pairs where they occurred), and then those within pair averages were averaged to give an overall average IPSC across all recorded pairs. This grand average was 105.5 ± 133.4 pA (mean ± standard deviation), with a median of 33.8 pA and a range of 7.1 to 417.1 pA (Figure 2). The amplitudes in individual pairs formed two separate clusters based on the size of the IPSC: a cluster with smaller IPSCs below 80 pA (n=23) and a cluster with a much larger IPSC between 220 and 420 pA (n=8). The average latency of the IPSC was 1.37 ± 0.47 ms, with a median of 1.27 ms and a range of 0.47 to 2.63 ms. No relation was apparent between the size of the IPSC and the latency (R^2^=0.01). Approximately half of the recorded pairs showed no failures of synaptic transmission across at least 25 consecutive trials (one presynaptic action potential = 1 trial; 15 out of 31 pairs). For those pairs that did show failures, the observed failure rates varied between 5.3% to 52.9%. Note that in this recording configuration, a ‘failure’ of transmission is only observed when all synapses making up that connection fail in that trial. This observed rate is thus, a composite failure rate. Overall, the composite failure rates were higher for the pairs with an IPSC<80 pA compared to pairs with IPSC>220pA (15.5% vs 2.7% average failure rate, p = 0.065), suggesting that pairs with smaller IPSCs might have fewer synapses.

**Figure 2.**
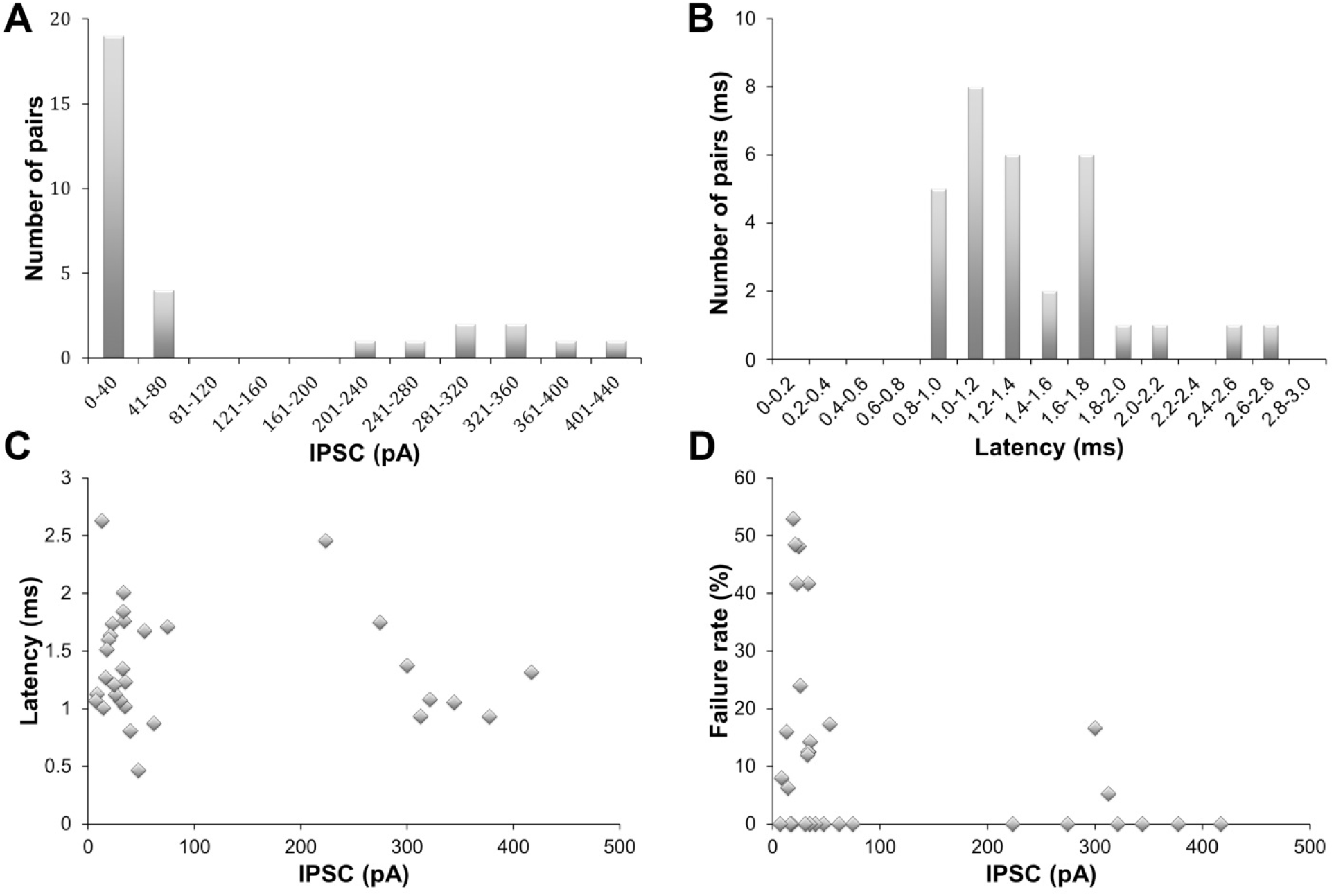
Electrophysiology of PV+ interneurons. **A.** Frequency distribution of the IPSC amplitudes of all pairs (n=31). **B.** Frequency distribution of the latencies of all pairs. **C.** Mean latencies plotted against mean IPSC. D. Failure rate against mean IPSC.

### Array tomography analysis - contact detection

Fourteen of the pairs with complete and bright neuronal fills that allowed tracing of distal dendrites and axons were processed for AT analysis. Based on the electrophysiological measurements, this sample was representative of the whole. The average evoked IPSC across these 14 pairs was 143.8 ± 144.7 pA (st. dev.), with a median of 48.8 pA and a range of 12.9 to 417.1 pA. The average latency of the response was 1.48 ± 0.55 ms (st.dev.), with a median of 1.33 ms and a range of 0.87 to 2.63 ms. Subsequent to tissue embedding and sectioning into 100 nM serial arrays, Neurobiotin-filled presynaptic interneurons were labeled on sections using Streptavidin Alexa 594, and Lucifer Yellow-filled postsynaptic neurons were immunostained using an anti-Lucifer Yellow antibody and a secondary Alexa 488 antibody (Figure 3). This resulted in a bright label of all neuronal domains, including the axon, which could be followed for long distances in these 14 well-filled samples. Because PV+ interneurons are known to preferentially target somata and proximal dendrites, a volume of 400 μm by 140 μm by 26 - 78 μm (H, W, D) centered on the postsynaptic neuron was examined for contacts between the two neurons. Contacts were identified as close appositions of a presynaptic varicosity (labeled with Streptavidin Alexa 594) and the postsynaptic neuron (immunostained with Lucifer Yellow antibody), that could be seen on at least 2 consecutive sections. Altogether, 131 contacts between the pre- and the postsynaptic neuron were found in the analyzed neuronal pairs. The synaptic nature of 35 randomly selected contacts was further confirmed by the presence of synaptic markers, such as synapsin on the presynaptic side and gephyrin on the postsynaptic side (Figure 3C). All of the 35 axonal varicosities examined were immunopositive for synapsin, and 91% of the targets were gephyrin-immunopositive, which is in accordance with previous studies showing that gephyrin is not detectable at all inhibitory synapses in mouse cortex (Collman et al. 2015).

**Figure 3.**
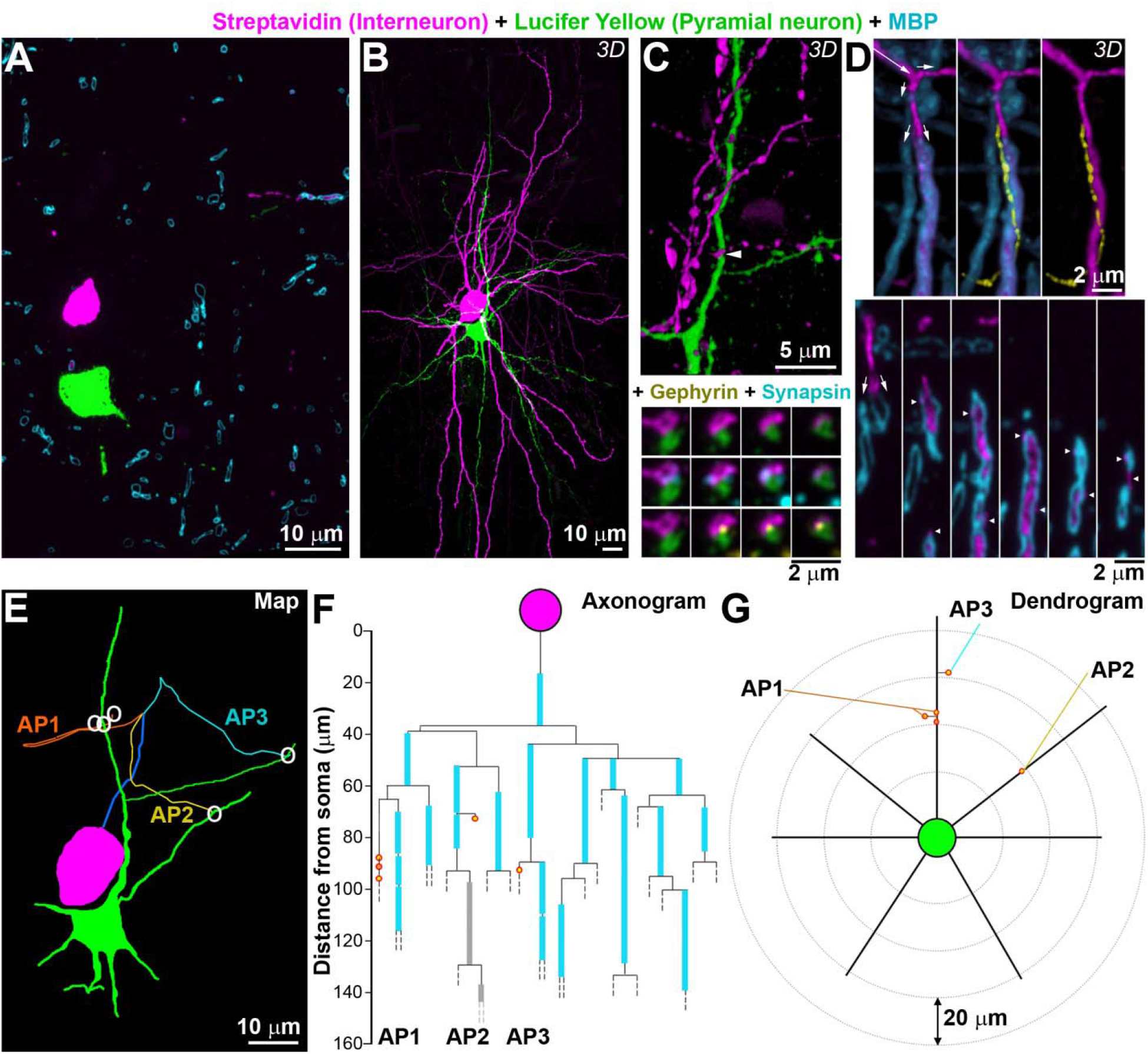
Neuronal pair mapping. **A.** Single section (100 nm thick) from pair MK181212, labeled with a Lucifer Yellow antibody (green, postsynaptic neuron), Streptavidin-Alexa594 (magenta, presynaptic interneuron) and MBP (cyan). **B.** MAX projection from the 727 sections through the same neuronal pair as A. C. Volume reconstruction of a contact between the presynaptic axon and the postsynaptic dendrite (top, arrowhead), with serial sections through it (bottom) labeled with synapsin (cyan) and gephyrin (yellow). D. (Top) Volume reconstruction of an axonal segment, three different views. Long arrow indicates the direction of the axon, and small arrows – bifurcations. At the 2 ^nd^ bifurcation, the axon divides into 2 branches, one myelinated and one unmyelinated. The unmyelinated axonal branch is shown in yellow in the middle and right panel. (Bottom) Selected single sections through the volume, small arrows point to the bifurcation, and arrowheads point the unmyelinated branch profiles. E. Map of the neuronal pair, with contacts represented by white circles. Each axonal path (AP) is color-coded. The beginning common axonal path (axon initial segment and first internode) is in blue. F. Axonogram of the interneuron in the same pair, with myelin in cyan, and partial (incomplete) myelination in grey. Contacts are shown as red circles, filled with orange (on dendrites) or yellow (spines). Dashed lines mark where the axons exited the traced volume. G. Dendrogram of the postsynaptic neuron.

### Axonal path tracing

From each identified synaptic contact, the axon was traced back to the presynaptic cell body (Figure3E). One of the pairs (MK180817-1) was excluded from this analysis because the two neurons were too far apart along the z-axis to follow the axons reliably across multiple sections. In addition, it was not possible to follow the axons for 10 synapses, usually due to axons exiting the imaged volume, or lost sections. The remaining 107 synapses were all traced back to the presynaptic cell body and the axonal path length was measured.

Myelinated internodes were identified by the presence of MBP immunolabel. Tracing of the axonal paths across serial ultrathin sections with immunofluorescent AT revealed features that would be very hard to detect using other light microscopy methods such as confocal microscopy. Because AT uses ultrathin sections, there is no out of focus fluorescence to increase background noise and obscure smaller features, and the z-resolution is effectively the thickness of the section, 100 nm in the present study, compared to 500 nm at best for confocal microscopy (Pawley 1995). Using AT we observed that unmyelinated axons branched extensively and were often characterized by very thin stretches connecting larger varicosities. Thin unmyelinated axons also sometimes ran immediately adjacent to larger myelinated axonal branches and ‘hid’ behind them (Figure 3D). The fact that array tomograms are digital reconstructions of very thin serial sections, allows the investigator to ‘see behind’ the obstruction to detect these ‘hiding’ axons.

Results from the axonal path tracing for each pair are summarized in Axonograms (Figure 3F) and the position of contacts on the postsynaptic cell are presented in Dendrograms (dendrogram representation inspired by Aliaga Maraver et al. 2018, Figure 3G). The axonograms and dendrograms for all pairs can be found in Supplemental Data.

### Basket cell axons are myelinated with a short AIS and short first internode

Several typical features emerged from the analyzed interneurons. In all but 2 of the interneurons, the axon originated from the apical part of the neuron, then ran up towards the pial surface before looping back and dividing into several branches to target the postsynaptic neuron. In many cases this visually resembled a ‘fountain’ (Figure 4A,B). Typically, the axon bifurcated after the first internode, then after a short distance each branch split in 2 or more branches. In two of the pairs, the axon split into branches before the first internode.

**Figure 4.**
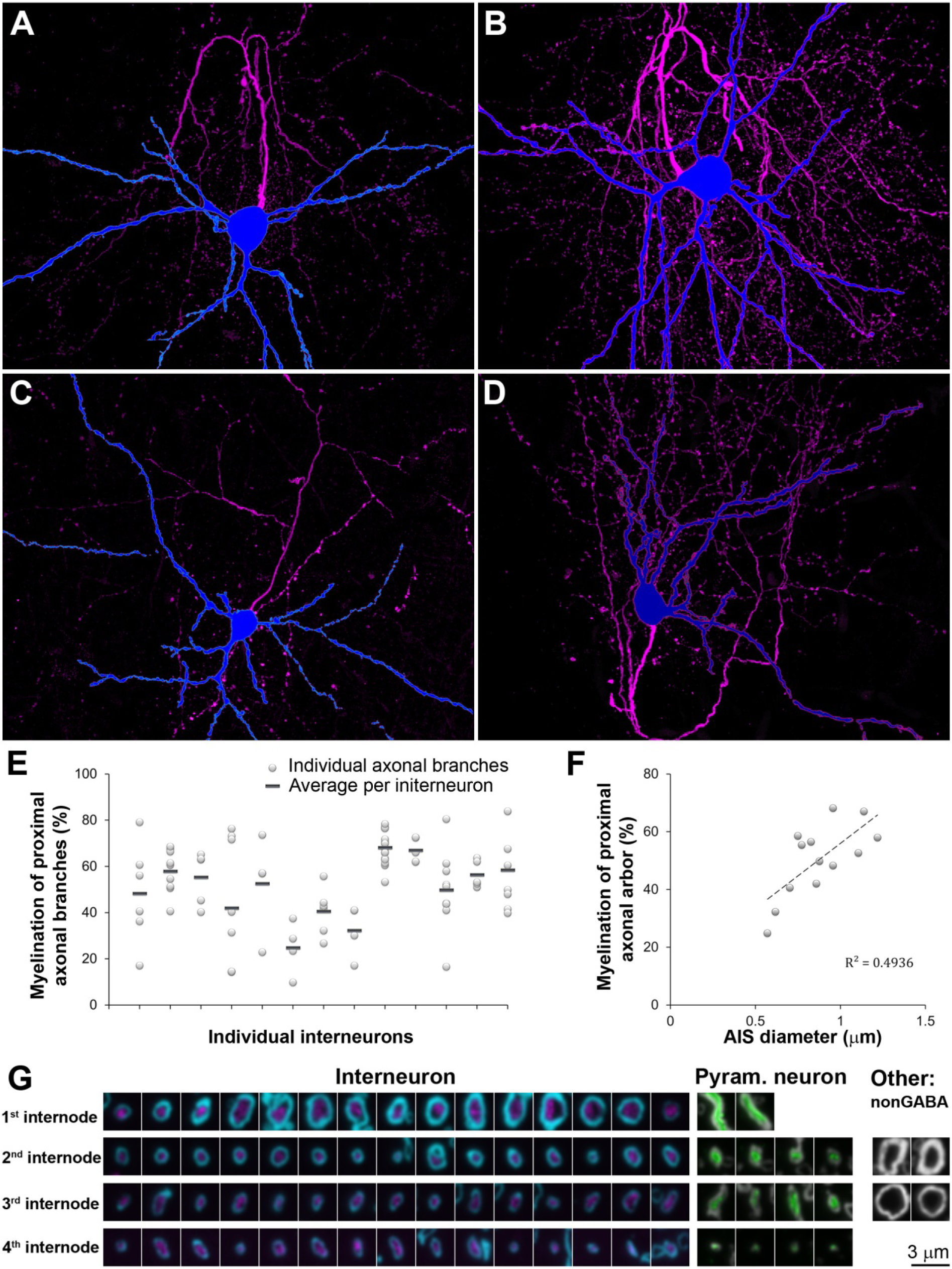
Features of interneuron axons and their myelin. A-D. Four interneurons (MAX projections through all sections) with the soma and dendrites in blue and axon in magenta. A: MK181221-1, B: MK190227, C: MK180814, D: MK181213. E. Percent length of the proximal axonal arbor (within 100 μm from the cell body) ensheathed by myelin. F. Relationship between the myelination of the proximal axonal arbor and the diameter of the AIS, p=0.125. G. Single sections through myelinated axons of interneurons (magenta, MBP in cyan), pyramidal neurons (green, MBP is grey), and unidentified nonGABA cortical axons (MBP in grey).

All of the interneurons were myelinated, consistent with previous studies of cortical PV+ basket cells (Micheva et al. 2016; Stedehouder et al. 2017). The length of the AIS (defined as the axonal stretch between its origin and the beginning of myelination) was remarkably similar regardless of cortical layer (Table 3). The average AIS length was 22.4 ± 2.7 μm (average ± st.dev), with a median of 22.4 μm, range 17.9 – 26.0 μm). This was in contrast to the large variation in AIS length for the four pyramidal neurons that had their myelinated axons within the reconstructed volumes: 24.7 μm, 25.0 μm, 65.0 μm, 101.4 μm). The diameter of the AIS was 0.87 ± 0.19 μm (mean ± st.dev), with a median of 0.87 μm, range 0.57 – 1.22 μm. This was substantially thicker than the diameter of the four pyramidal neurons’ AIS (0.52 μm, 0.59 μm, 0.59 μm, 0.65 μm).

**Table 3.**
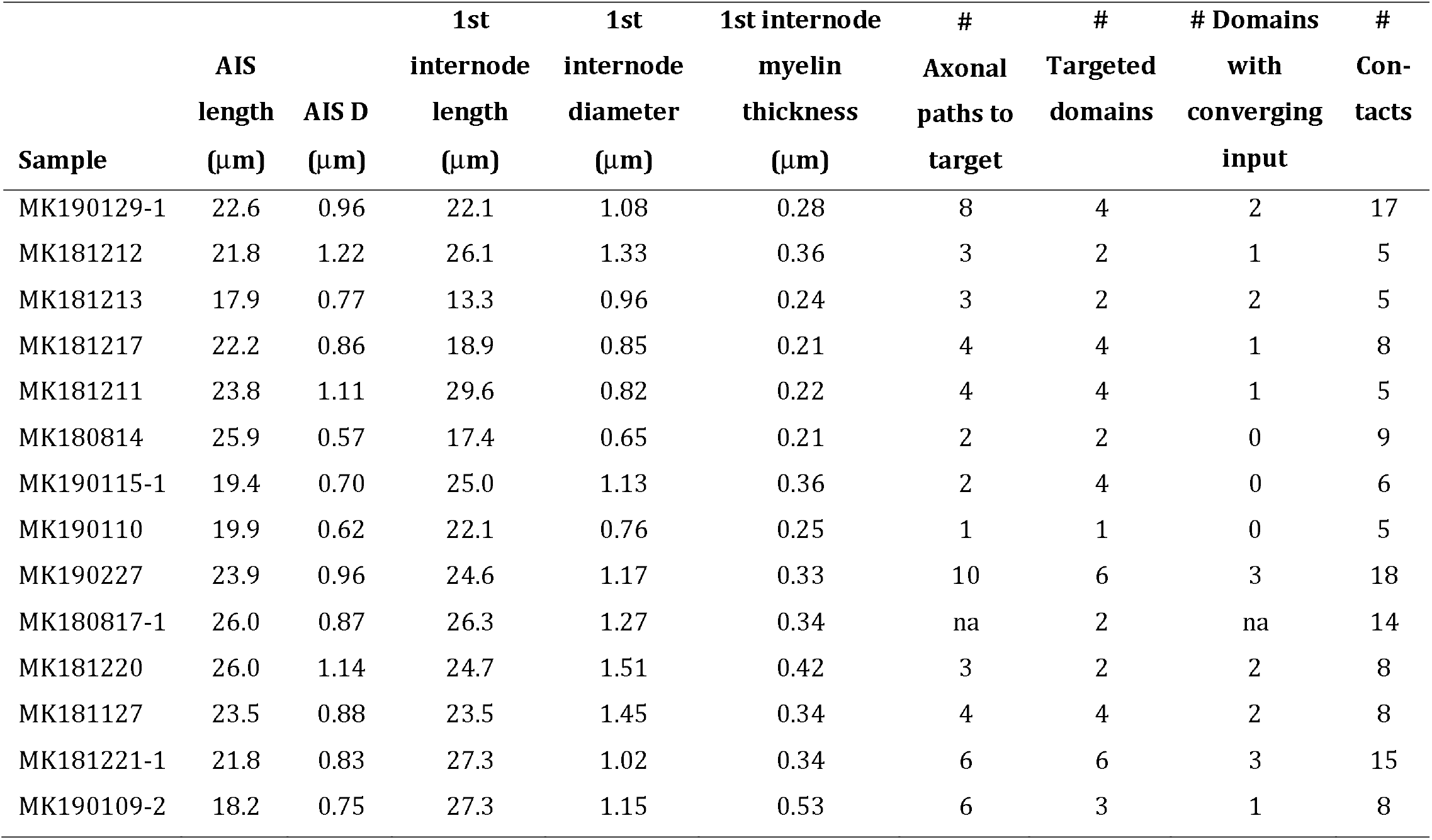
Morphological measurements of individual connections.

|  | AIS |  | 1st internode | 1st internode | 1st internode | # | # | # Domains | # |
| --- | --- | --- | --- | --- | --- | --- | --- | --- | --- |
|  | length | AIS D | length | diameter | myelin thickness | Axonal paths to target | Targeted domains | with converging input | Con-tacts |
| Sample | ($\mu\text{m}$) | ($\mu\text{m}$) | ($\mu\text{m}$) | ($\mu\text{m}$) | ($\mu\text{m}$) | | | | |
| MK190129-1 | 22.6 | 0.96 | 22.1 | 1.08 | 0.28 | 8 | 4 | 2 | 17 |
| MK181212 | 21.8 | 1.22 | 26.1 | 1.33 | 0.36 | 3 | 2 | 1 | 5 |
| MK181213 | 17.9 | 0.77 | 13.3 | 0.96 | 0.24 | 3 | 2 | 2 | 5 |
| MK181217 | 22.2 | 0.86 | 18.9 | 0.85 | 0.21 | 4 | 4 | 1 | 8 |
| MK181211 | 23.8 | 1.11 | 29.6 | 0.82 | 0.22 | 4 | 4 | 1 | 5 |
| MK180814 | 25.9 | 0.57 | 17.4 | 0.65 | 0.21 | 2 | 2 | 0 | 9 |
| MK190115-1 | 19.4 | 0.70 | 25.0 | 1.13 | 0.36 | 2 | 4 | 0 | 6 |
| MK190110 | 19.9 | 0.62 | 22.1 | 0.76 | 0.25 | 1 | 1 | 0 | 5 |
| MK190227 | 23.9 | 0.96 | 24.6 | 1.17 | 0.33 | 10 | 6 | 3 | 18 |
| MK180817-1 | 26.0 | 0.87 | 26.3 | 1.27 | 0.34 | na | 2 | na | 14 |
| MK181220 | 26.0 | 1.14 | 24.7 | 1.51 | 0.42 | 3 | 2 | 2 | 8 |
| MK181127 | 23.5 | 0.88 | 23.5 | 1.45 | 0.34 | 4 | 4 | 2 | 8 |
| MK181221-1 | 21.8 | 0.83 | 27.3 | 1.02 | 0.34 | 6 | 6 | 3 | 15 |
| MK190109-2 | 18.2 | 0.75 | 27.3 | 1.15 | 0.53 | 6 | 3 | 1 | 8 |

In all of the myelinated interneurons, myelin ensheathed only the proximal part of the axonal arbor, and myelinated internodes were often interspersed with longer unmyelinated stretches of axons. There were large individual variations in myelination, both between interneurons, and between axonal paths of the same interneuron (Figure 4E). The degree of proximal axonal myelination (% myelinated axonal length within 100 μm from the cell body) correlated well with the thickness of the axon initial segment (R=0.71, p < 0.01 Figure 4F), as well as with the thickness of the 1^st^ and 2^nd^ internodes (R=0.70 and R=0.74 respectively, p < 0.01).

The first internode of the basket cells was short, with a length of 23.4 ±4.4 μm (average ± st.dev), with a median of 24.6 μm and range 13.3 – 29.6 μm. The first internode diameter was 1.08 ± 0.26 μm (average ± st.dev), with a median of 1.10 μm, range 0.65 – 1.51 μm). The diameter of the myelinated internodes of the basket cells was substantially larger than the diameter of the myelinated internodes of the pyramidal neurons (first internode diameter 0.54 ± 0.07 μm, median of 0.57, p < 0.01; Figure 4G). There were, however, other very large diameter axons of presumably excitatory neurons that were GABA immunonegative and did not belong to the filled neurons, indicating that in neocortex, the large diameter myelinated axons are not an exclusive feature of basket cells.

### Synaptic contacts cluster close to postsynaptic soma

Each interneuron made on average 9.4 ± 4.7 contacts on the postsynaptic neuron, with a median of 8 and range of 5 to 18. Overall, among the 131 contacts identified, 25.2% were on cell bodies, 73.3% were on dendrites, and 1.5% were on axon initial segments, rather similar to what was previously reported when analyzing targets of basket cell axons using large-scale serial section electron microscopy (12.6 % on cell bodies, 85.5 % on dendrites and 1.9 % on axons; Micheva et al., 2016). The contacts targeting dendrites tended to be close to the cell body, with an average distance of 37.6 ± 26.6 μm from the edge of the cell body (median of 30.6 μm, range 1– 117 μm, Figure 5B). When the postsynaptic target was an interneuron, as many as 36% of the contacts were on the somata, compared to 23% on a postsynaptic pyramidal neuron (Figure 5A). The dendritic targets on interneurons were also much closer to the soma, on average 18.1 ± 10.3 μm vs. 41.5 ± 27.2 μm away from the soma when the postsynaptic cell was a pyramidal neuron (p<0.01, Figure 5B).

**Figure 5.**
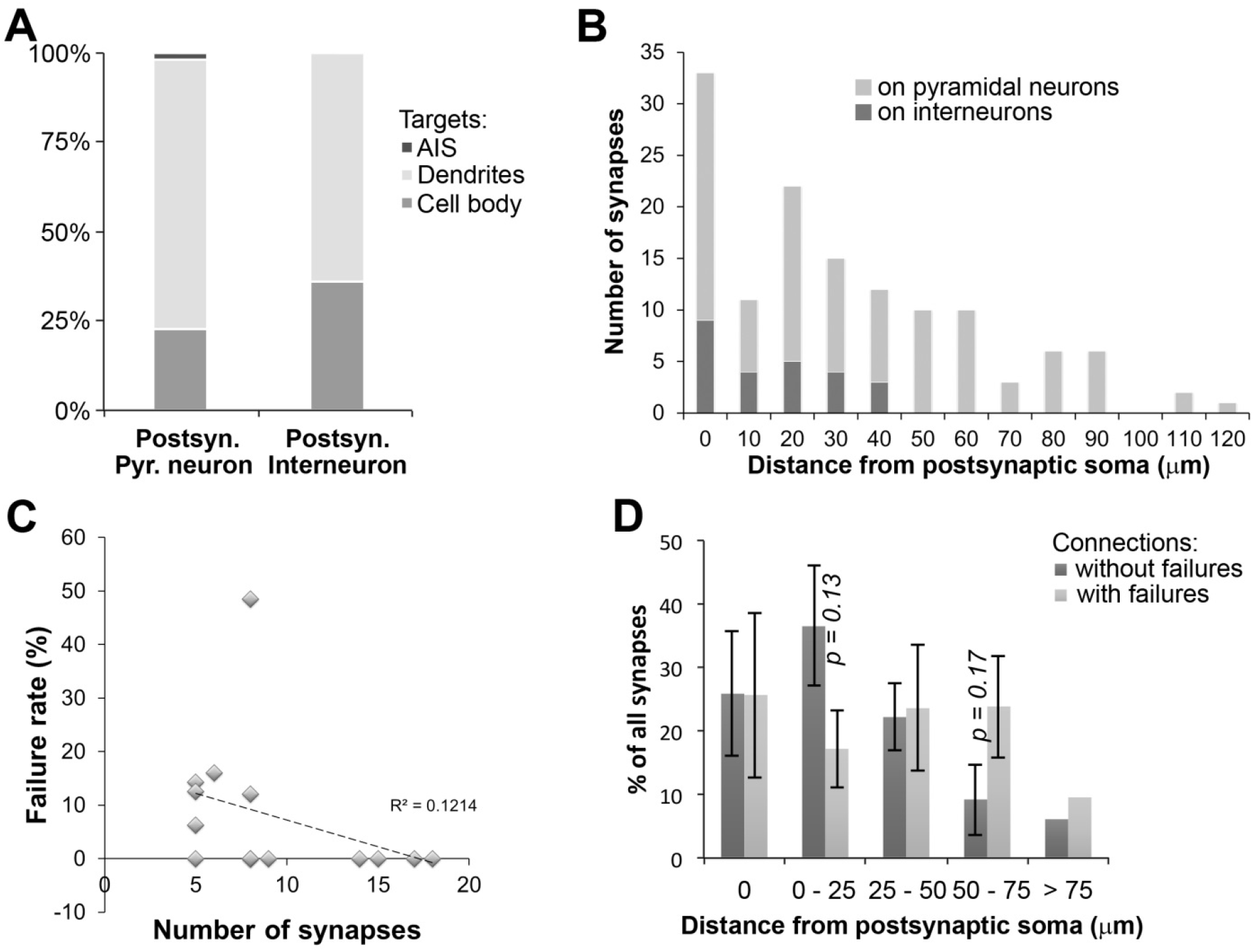
Postsynaptic targets of interneurons and their relation to failure rates of the connection. A. Postsynaptic targets of interneurons, depending on the postsynaptic neuron cell type. B. Frequency distribution of the distances of each synapse from the postsynaptic cell body in the analyzed pairs (n=14). C. Failure rate plotted against the number of contacts in a connection. D. Comparison of synapse distribution on the postsynaptic neuron between connections without and with failures in neurotransmission.

As expected, the recorded current in the postsynaptic neuron related to the number of synaptic contacts in a pair. Thus, pairs with large IPSC>220pA had significantly more synapses compared to pairs with IPSC<80pA (6.3 ± 1.5 vs 13.5 ± 4.1, p<0.001, n=8 and n=6 respectively).

### Connection failure rates and synapse release probabilities of PV+ interneurons are highly variable

The analysis of all recorded pairs showed that pairs with large IPSC>220pA, had very few composite failures of synaptic transmission (Figure 2D). We make the distinction of ‘composite’ failure because each pair of cells has multiple synapses made between them, and thus what we observe as a ‘failure’ of transmission in response to a presynaptic action potential, is actually a failure of all synapses that make up that connection in that trial. If even a single synapse, at least a synapse terminating electrically close to the soma, successfully transmits in response to a presynaptic action potential, we would record that as a ‘success’. Because all the neuron pairs we analyzed have multiple synaptic contacts, we hypothesized that this composite failure rate would be higher in connections with fewer synapses. However, we found only a very weak negative correlation between the number of synapses and the failure rates of a connection (R^2^ = 0.12, Figure 5C). For example, in the 4 connections with 8 synapses each, the composite failure rates ranged between 0 to 48%. We therefore explored other factors that might influence the reliability of synaptic transmission between the PV+ interneuron and its postsynaptic neuron, such as position of the synapses relative to the postsynaptic cell body or release probability at individual synapses. Comparing the positions of synapses on the postsynaptic neuron, the pairs with no detected failures had the same proportion of synapses targeting the soma as the pairs with failures (26%), but they had about twice higher proportion of synapses within 25 μm of the cell body (37 vs 17%). This difference, however, was not statistically significant (p=0.13) (Figure 5D).

Because for the pairs analyzed with AT both the number of synaptic contacts and the failure rates of the connections were known, we used this to estimate the release probability of individual synapses. For these calculations we assumed that all the synapses in a pair have similar release probabilities, which is not necessarily true (Branco and Staras 2009). The average release probability of a PV+ interneuron individual synapse calculated from the 6 connections with failures was estimated at 0.28 ± 0.12, with a median of 0.29 and range of 0.09 to 0.43. These estimates of release probability underscore the large variability of PV+ interneurons synaptic function.

### Basket cells contact the postsynaptic cell via multiple axonal paths with different geometries and myelination

In all but one of the 13 pairs with reconstructed axons, the interneuron contacted its postsynaptic target by several different axonal paths (Table 3, Figure 6). After the first branching point there were typically two axonal paths synapsing onto the postsynaptic neuron (average of 1.85 ± 0.55, median of 2), and after the 2^nd^ branching point −3 (average of 2.54 ± 0.88, median of 2). Only in one pair (MK190110), the presynaptic interneuron targeted the postsynaptic cell via a single axonal branch. The number of axonal paths (separated by 2 or more branching points) correlated very well with the number of contacts (R^2^=0.78), with an average of 1 path per 2 contacts.

**Figure 6.**
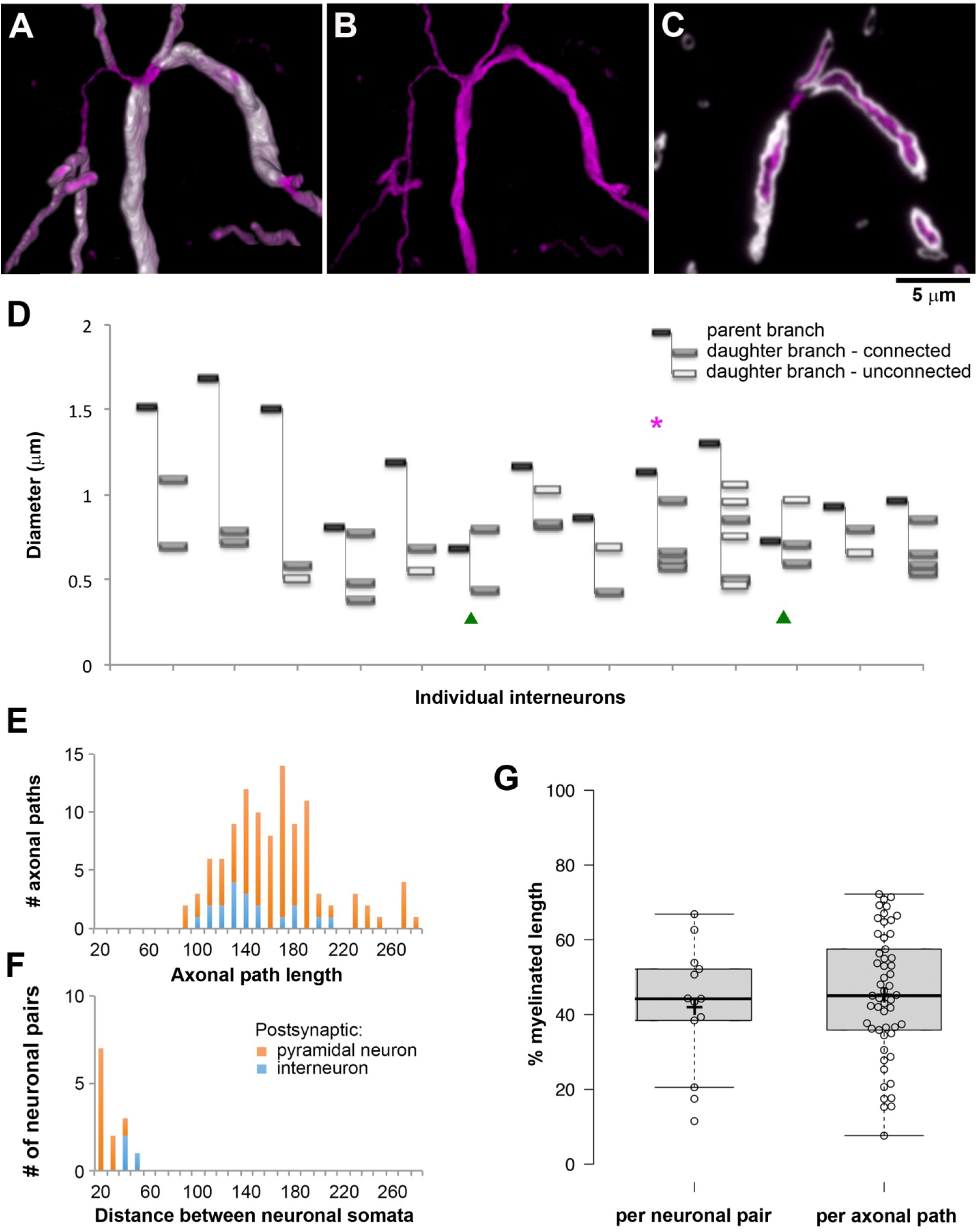
Axonal branching. **A.** The primary axon of the interneuron from pair MK190227 (Figure 3B) divides into 4 daughter branches with varying diameter. Volume reconstruction; interneuron axon labeled with Streptavidin-Alexa594 (magenta), myelin with MBP (white). **B.** Same as A, without the myelin immunolabel. **C.** Single section (100 nm) through the same axon as A and B. D. Thickness of individual axons at the first branchpoint. Magenta asterisk points to the interneuron from panels A-C. Green arrowheads identify the interneurons where the first branchpoint occurred immediately after the axon initial segment. In all the other interneurons the axon divided after the first internode. E. Frequency distribution of the lengths of the axonal paths to the postsynaptic contacts in the pairs, color-coded depending on the postsynaptic cell type. **F.** Frequency distribution of the distance between the neuronal cell bodies in the pairs. G. Boxplots of the percent myelinated length of the axonal paths, presented as an average for each pair (n=13, left) and per individual axonal path (n=53, right). Center lines show the medians; box limits indicate the 25^th^ and 75^th^ percentiles as determined by R software; whiskers extend 1.5 times the interquartile range from the 25^th^ to the 75^th^ percentiles; crosses represent sample means; data points are plotted as open circles.

The individual axonal paths for the same neuronal pair varied in their thickness. This was particularly obvious at axonal branching points. In the example presented in Figure 6, the axon divides into 4 branches that all form contacts with the postsynaptic neuron. The diameter of one of these branches is almost twice as large as the diameters of the other three branches (0.96 μm, vs. 0.57, 0.60 and 0.66 μm), and it is also ensheathed by thicker myelin (0.58 μm, vs. 0.46 μm for two of the other branches, and the fourth branch is unmyelinated until it divides into 3 myelinated branches).

All imaged pairs consisted of neurons that were either immediately adjacent or very close by, with a distance between their cell body centers of 16 to 40 μm (25 ± 9 μm, average ± st.dev.; 19.5 μm median; Figure 6F). Nonetheless, the lengths of the axonal paths connecting the two neurons were much longer, 159 ± 41 μm (average ± st.dev.), with a median of 154, and a range of 88 to 272 μm; Figure 6E). The great majority of axonal paths (83%) were between 100 and 200 μm long. The postsynaptic target neuron was a pyramidal neuron in most of the pairs, except for 3 pairs where it was an interneuron. Even though the distance between the neurons in these 3 pairs was significantly larger (36 ± 4 μm vs. 22 ± 8 μm, p=0.01), the axonal paths connecting interneurons were shorter than the axonal paths connecting a PV+ interneuron and a pyramidal neuron (139 ±31 μm vs. 163 ± 42 μm, p=0.02).

All of the individual axonal paths in the imaged pairs were myelinated, but to a varying extent (Figure 6G), with an average of 45 ± 17%, median of 45% and range of 8 to 72% of the axonal path length ensheathed by myelin. On average there were 3 ± 1 internodes per axonal path, median of 3, range 1 – 6. There were no differences between the axonal paths connecting a PV+ interneuron and a pyramidal neuron vs. PV+ interneuron and an interneuron. The extent of myelination was not correlated with the length of the axonal path (R2=0.0002).

### Conduction velocity is higher along more heavily myelinated axons

While it is well known that myelination increases the speed of action potential propagation in long-distance projecting axons (Waxman and Bennett 1972; Moore et al. 1978), the contribution of myelination to the timing of signal propagation along local axons of PV+ interneurons is much less understood. To address this question we estimated the conduction velocity along PV+ interneuron axons by dividing the length of the shortest axonal path in a connected pair by the latency of response measured during the paired electrophysiological recordings (Table 4). Axonal paths to distal targets (> 50 μm from the postsynaptic cell body) were not included as they were unlikely to contribute to the IPSC recorded at the soma (Kubota et al. 2015). The estimated conduction velocity ranged between 0.03 m/s to 0.13 m/s and correlated well with the extent of myelination of the axonal paths (R=0.64, p=0.02; Figure 7A). This estimate for conduction velocity does not take into account the synaptic delay, i.e. the time between the arrival of the action potential at the synaptic contact and the postsynaptic response, and is therefore an underestimate. A value of 0.5 ms is often used as an estimate of this synaptic delay (Katz and Miledi 1965). However, because some of our recorded pairs had an IPSC latency (conduction time + synaptic delay) of less than 0.5ms, we surmised that the delay in these synapses is less. For example, the average latency in our fastest recorded pair was 0.47 ms (range of 0.3 – 0.7 ms), suggesting that synaptic delay must be significantly less than 0.3 ms. To be conservative, we thus used an estimate of 0.25 ms for synaptic delay, consistent with the very fast signaling properties of PV+ interneurons (Hu et al., 2014) and less than the often quoted value for synaptic delay of 0.5 ms. Assuming a synaptic delay of 0.25 ms would result in conduction velocities between 0.04 and 0.17 m/s with an average of 0.13 ± 0.04 ms, that are again well correlated with the degree of myelination (R=0.68, p=0.01; Figure 7B).

**Figure 7.**
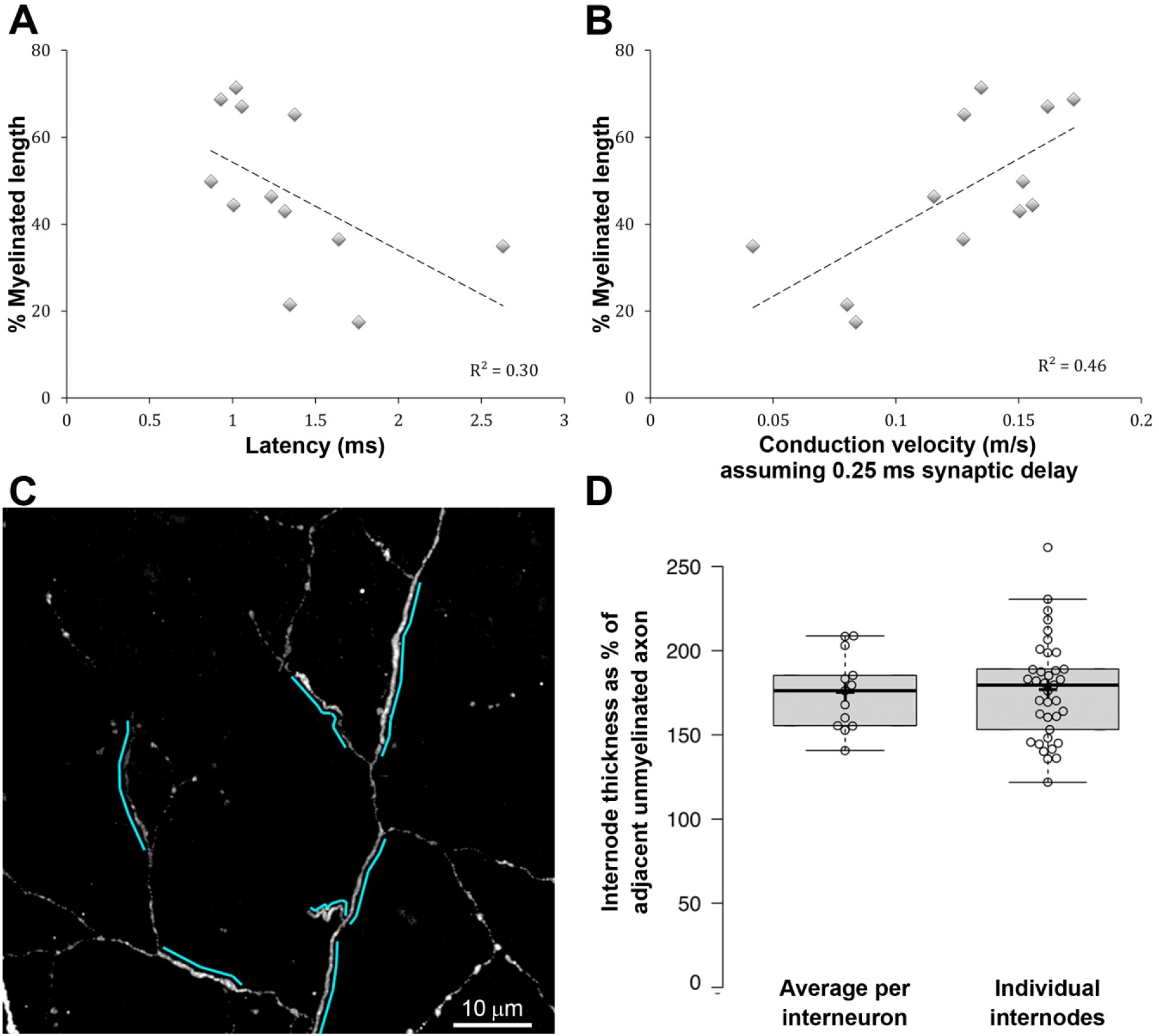
Conduction velocity varies with myelination. **A.** Correlation between latency and the percent length of the shortest axonal path ensheathed by myelin, p = 0.065. B. Correlation between conduction velocity and the percent length of the shortest axonal path ensheathed by myelin, accounting for a synaptic delay of 0.25 ms, p = 0.01. C. MAX projection featuring the axon of interneuron MK180814. Myelinated stretches of the axon, indicated by a cyan line, are visibly thicker than the adjacent unmyelinated axon. **D.** Diameter of axons at 2^nd^ and 3^rd^ internodes as a percent of the diameter of the adjacent unmyelinated axon.

**Table 4.**
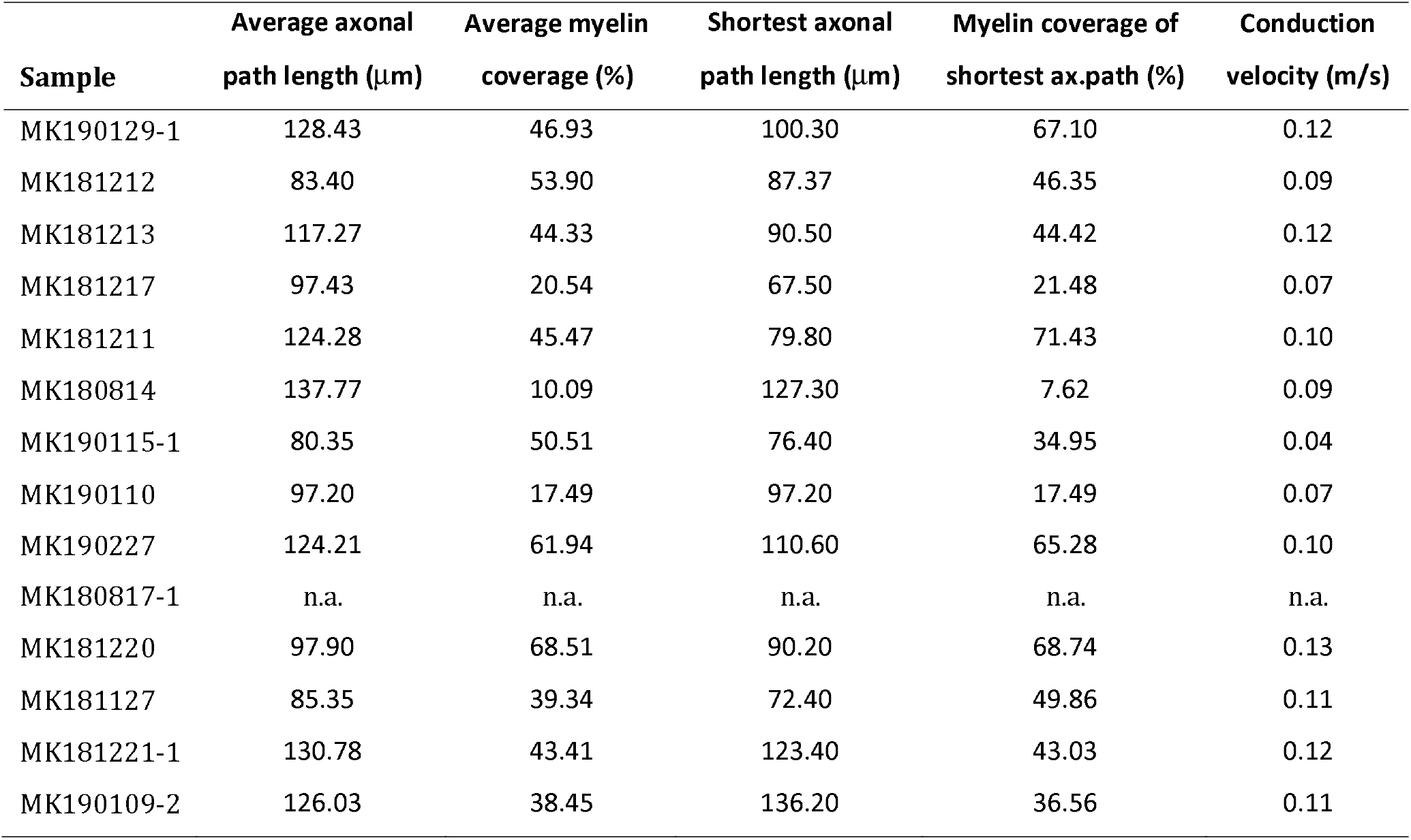
Axonal path lengths, myelination and conduction velocity.

In addition to the faster saltatory conduction in the presence of myelin, another variable that affects conduction velocity is axon diameter, with larger axons having faster propagation of action potentials (Hodgkin and Huxley 1952). Interestingly, the myelinated axonal internodes in our sample consistently appeared to have larger diameters compared to the adjacent unmyelinated axonal segments. The only exception was the first internode, which was thinner than the preceding axon initial segment. To quantify the thickness of myelinated vs. unmyelinated axonal segments, we measured the axonal diameter in secondary and tertiary internodes and in the adjacent unmyelinated segments before and after the internode. Invariably, the myelinated internodes had a significantly larger diameter than the adjacent unmyelinated segments of the axon (0.83 ± 0.17 μm vs. 0.48 ± 0.07 μm, respectively, p < 0.001), (Figure 7B,C), consistent with previous observations (Stedehouder et al. 2019). Thus, the high conduction velocity along the axons of neocortical PV+ interneurons is likely assisted by at least two interrelated mechanisms: myelination and an increase in axon diameter.

### Conduction velocity is higher at connections with different axonal paths converging onto the same domain of the postsynaptic neuron

As described above, the interneuron generally contacted its postsynaptic target by several different axonal paths (Table 3 and Figure 6). This ‘redundancy’ of connections was apparent at the postsynaptic cell too, as the postsynaptic neuron received basket cell input on multiple domains (Figure 8). A cell domain was defined as either the cell body, or a primary dendrite with its branches. In cases were the primary dendrite branched into two large branches close to the cell body (within 20 μm), these two branches were considered as separate domains. On average, 4 different domains of the postsynaptic neuron were contacted in each pair. There appeared to be no preference for specific domains in a connection, for example apical or basal dendrites, unlike what was reported in some cases (Thomson et al. 1996). In 3 of the pairs, including the one shown in Figure 8A, the postsynaptic pyramidal neuron received input from the interneuron on its apical dendrite, basal dendrites and soma (and axon initial segment in one of these pairs).

**Figure 8.**
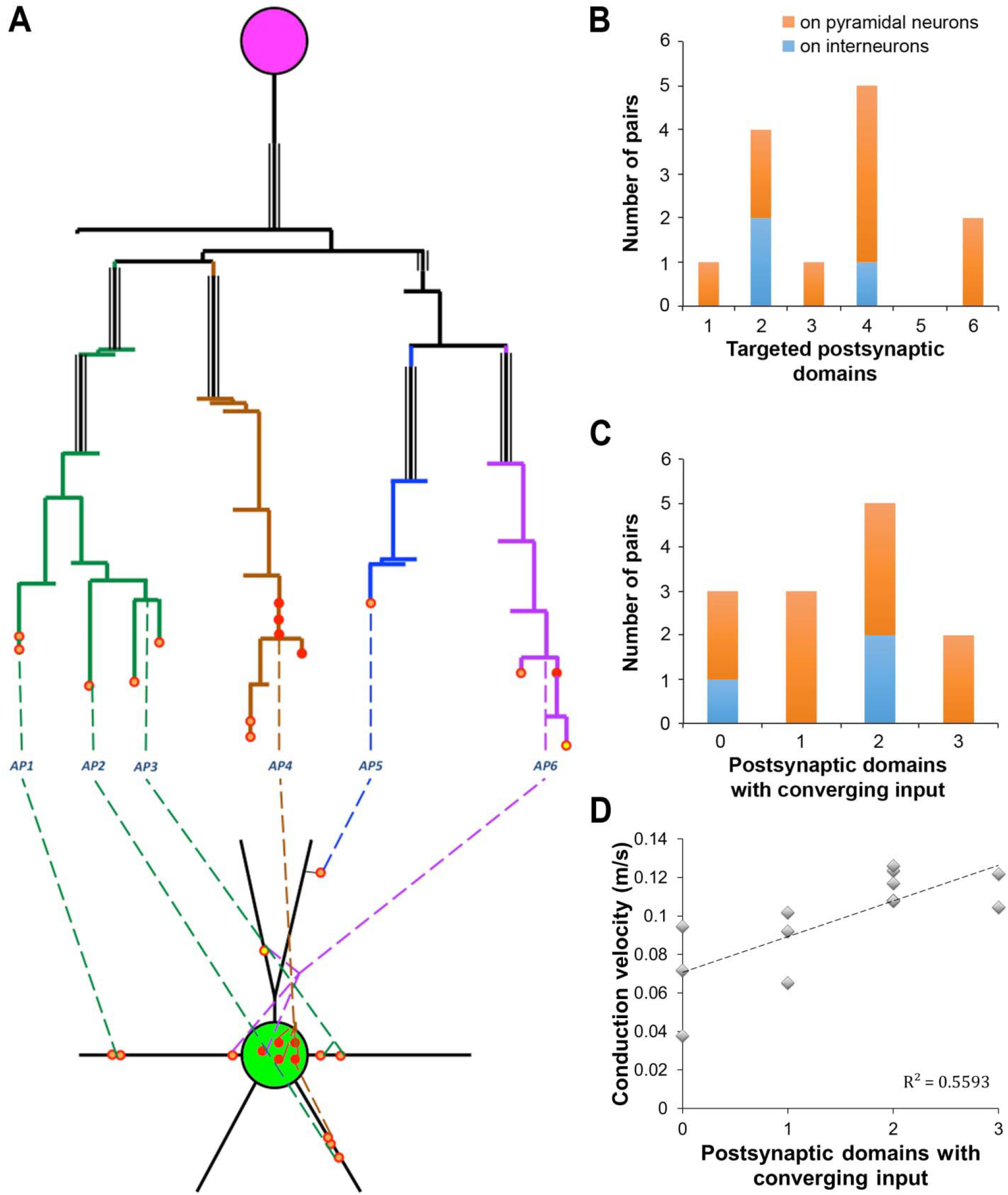
Multiple axonal paths and targets within a neuronal pair. **A.** Map of neuronal pair MK181221-1, with the interneuron making 15 contacts onto the postsynaptic pyramidal neuron. The numbered axonal paths here are defined as those that are separated by two or more branching points. Axonal paths with the same color originate from the same 3 ^rd^ or 4^th^ order axonal branch. The same axonal paths (e.g. AP4 and 6) can contact more than one postsynaptic domain, and the same postsynaptic domain can receive input from more than one axonal path (e.g. the postsynaptic cell body and the one of the basal dendrites). **B.** Distribution of the number of targeted postsynaptic domains per neuron among the mapped pairs. Bars are color-coded depending on the postsynaptic neuron. C. Distribution of the number of postsynaptic domains with converging input per neuron. Bars are color-coded depending on the postsynaptic neuron. For B and C, n=10 PV+ interneuron – pyramidal neuron pairs, and n=3 PV+ interneuron - interneuron pairs. D. Correlation between conduction velocity and the number of postsynaptic domains with converging input from different axonal paths, p < 0.01.

Almost half of the contacted postsynaptic domains (43%) received converging input from 2 or more presynaptic axonal paths (Figure 8C,D). For example, the cell body of the pyramidal neuron in pair MK181221-1 (Figure 8A) received input from 2 different axonal paths, and so did one of its basal dendrites, where the 2 axonal paths terminated very close by, within less than 5 μm. Surprisingly, there was a strong and significant correlation between the number of postsynaptic domains with converging input and the conduction velocity (R = 0.75, p < 0.01). While it is unclear if any causation exists between these two variables, it is reasonable to expect that the convergence of different axonal paths onto the postsynaptic target will enhance the reliability of the connection and safeguard against potential branch failures. Thus, the mesh-like connectivity between the basket cell and its postsynaptic target may be one more specialization of these neurons, along with myelination and many others, that ensures fast and efficient neurotransmission.

## Discussion

The present study was motivated by the question whether myelin can contribute in any biologically meaningful way to the speed of action potential propagation along the local axons of PV+ basket cells. This question arises because interneurons project locally, with generally short axons. Is there enough axon length to allow any conduction speed conferred by myelin to be of sufficient magnitude to make even a detectable difference in axon conduction velocity? We sought to answer this question by making electrophysiological recordings from pairs of neurons consisting of a presynaptic PV+ basket cell, and a single postsynaptic target neuron; usually a pyramidal neuron, but sometimes another interneuron. We evoked single action potentials via current injection into presynaptic interneuron and then measured the amplitude, latency and failure rate of inhibitory postsynaptic currents (IPSCs) that resulted in the postsynaptic neuron. We subsequently used Array Tomography (AT) of those same neuron pairs to precisely map their connectivity, including axon length and the extent of myelination. Knowing the length of the axon path between the pre- and postsynaptic cell, and the previously recorded latency, we could estimate a conduction velocity, and further examine relationships between the characteristics of the axon path(s) and the characteristics of the evoked IPSC.

We show that the extent of myelin coverage varies widely between PV+ neurons and significantly correlates with the conduction velocity along these axons. This is consistent with the well-known role of myelin in speeding up neurotransmission along long-range projections. Even at the short rage of the local PV+ basket neuron connections, 100 – 200 μm, variations in myelin coverage can introduce a difference in the latency of synaptic transmission of 1 ms or even more (Figure 7A), which could have important functional consequences, for example by affecting the magnitude of coordinated oscillations (Pajevic et al. 2014) that are controlled by PV+ interneurons or by disrupting the high-precision feed-forward circuit that prevents the propagation of highly synchronous activity (Schmidt et al. 2017). Additionally, we reveal a unique connectivity pattern between individual PV+ basket cells and their postsynaptic partner, with multiple axonal paths contacting the postsynaptic neuron at multiple domains. While there are some examples of this in the literature (Thomson et al. 1996), our study highlights this feature of basket cell axonal organization as the rule, rather than an interesting exception. The different axonal paths by which the interneuron contacts the postsynaptic cell vary in axonal diameter, extent of myelination, and location of the postsynaptic targets. The possibility of differential activation of daughter branches at axonal branch points (Debanne 2004) would thus enable the interneuron to elicit a variety of postsynaptic responses, ranging from powerful inhibition of the postsynaptic neuron to selective inhibition of a specific input to individual dendritic branches or spines. In almost half of the pairs, different axonal paths converge onto the same postsynaptic domain, and this arrangement is characteristic of PV+ basket cell connections with higher conduction velocity. This is likely another specialization of PV+ basket cells that ensures reliable neurotransmission, guarding against potential branch failures.

The present results further confirm and complement the current understanding of the connectivity of PV+ basket cells which have been the focus of extensive studies over the past 35 years (Hu et al. 2014). Recent large-scale electron microscopy studies have revealed unprecedented details of the axonal organization (Schmidt et al. 2017) and the postsynaptic targets of cortical PV+ basket cells (Kubota et al. 2015). However, except for a few classical examples (Somogyi et al. 1983; Thomson et al. 1996; Tamas et al. 1997), not much information is available on the organization and function of individual basket cell-to-postsynaptic neuron connections which, given the variability of neuronal structure and function, is indispensable to understanding the rules of PV+ basket cell physiology. The paucity of data on individual pair connectivity is due to a great extent to the very high labor intensity of the usual technique of choice, electron microscopy, to image the large volumes needed to follow the highly branched and meandering axons of PV+ basket cells in order to capture all their connections with a postsynaptic target. Furthermore, combining electron microscopy with another laborious technique of dual whole cell recordings to identify synaptically connected neurons and obtain functional data from them, multiplies the complexity and time constraints of the task. In the present study, using high-resolution imaging with immunofluorescent AT instead of electron microscopy allowed us to considerably speed up this process, while still being able to resolve enough detail to follow the PV+ basket cell axons (e.g. Figure 3).

The axons of PV+ basket cells are significantly thicker than the axons of pyramidal neurons (Figure 4, and also Schmidt et al. 2017), and their synaptic boutons are among the larger ones in cortex with a diameter of 1 to 2 μm (Thomson et al. 1996), which made them ideal targets for light-level AT detection and mapping. With our approach we confirmed a number of previous observations on PV+ basket cells and their axonal organization. Thus, all the basket cell axons in our sample were myelinated with a short AIS and short first internode. Only the proximal part of the axonal arbor (typically within 150 μm from cell body) was myelinated, and myelinated internodes were often interspersed with longer unmyelinated stretches of axons. The synaptic contacts made by these axons clustered close to the postsynaptic soma. Indeed, these features of basket cell connectivity appear highly stereotypical, as they are found in different species, and different cortical areas and layers. It is rather astonishing that the connectivity patterns first discerned from EM reconstructions of only 3 basket cells in layers ¾ of cat striate cortex (Somogyi et al. 1983) are so consistent with the basket cells in layers 3 through 6 of the perirhinal and entorhinal cortex of mice from the present study.

Several new observations on the axonal organization of PV+ basket neurons emerged from our study. For example, the length of the axon initial segment (AIS), identified as the stretch between the axon hillock and the beginning of the myelin of the first internode, is short and remarkably consistent (average of 22 μm with a range of only 18 to 26μm), which is very different from the highly variable AIS of pyramidal neurons that can extend from around 20μm to more than 100μm (Tomassy et al. 2014). Because the AIS is a specialized structure with very high density of voltage-gated ion channels and the site of action potential initiation (Huang and Rasband 2018), its uniform length is consistent with the reliability of PV+ basket cells firing. On the other hand, the AIS diameter is more variable and correlates with the degree of axonal myelination, suggesting a coordinated organization of the entire axonal arbor, in support of the emerging notion that myelinated axons represent functional units (Suminaite et al. 2019).

The axonal path length in the connected pairs in our sample is always much longer than the geometric distance between the two neurons, with the great majority of axonal paths (83%) falling between 100 and 200 μm, while the largest distance between the two neurons in our pairs was only 40 μm. Indeed, one interesting and characteristic feature of the axon arbors we have examined in this study is that they almost always exit the interneuron with the AIS oriented toward the cortical surface. Despite the fact that our chosen pre- and postsynaptic neurons were always close to each other, and generally in the same cortical layer, in all but one case the interneuron axon proceeded away from the target neuron, before taking a “u-turn” and heading back to its termination on the postsynaptic target neuron. This was interesting because this makes the axon longer than it ‘needs’ to be if the goal were simply to connect to the postsynaptic cell with the shortest possible axon. Might there be some reason why this system evolved this way? One idea is that neural rhythms generated by local feedback circuits, which are known to depend on interneuron activity, require some delay in the arrival of the inhibitory synaptic transmission, which is provided by a longer axon. In this context, we might better understand the patchy nature of the myelination of PV+ interneuron axons not so much as a mechanism for providing maximum speed (which would be aided by full myelination), but rather as a way of tuning the latency to optimize the neuron’s role in maintaining circuit rhythms. Furthermore, because the great majority of local connections of PV+ basket cells are within a radius of 200 μm (Packer and Yuste 2011), the 100 – 200 μm range of axonal path length may be an optimal target length to ensure synchronous communication with all the local postsynaptic targets. Even though our sample included only pairs of neurons that were within 40 μm or closer, it is likely that when the connected neurons are 100 to 200 μm apart the axonal paths may take a much more direct route, as evidenced by the fact that in the examined pairs in addition to the characteristic “u-turn”, other branches of the same axon proceed straight up into the upper cortical layers and presumably make contacts with neurons in those layers.

There are several caveats and limitations to this study. While the use of immunofluorescent AT to map PV+ basket cell connectivity highly increased the output of our study, the identification of synaptic contacts relied on structural features (presence of an axonal varicosity directly apposed to the postsynaptic neuron), and was additionally confirmed in about a quarter of the cases by further immunostaining with pre- and postsynaptic markers. It is therefore possible that some synaptic contacts were missed, however, this is likely a small minority, because the number of synaptic contacts that we identified is highly consistent with what is reported in the literature (Somogyi et al. 1983; Tamas et al. 1997). What was not possible with our approach, however, was to reliably distinguish between a synaptic contact on a dendritic shaft or on spine, or even, as often observed in EM studies (Somogyi et al. 1983), a contact on both a dendritic spine and adjacent dendritic shaft by the same presynaptic bouton. Therefore, for our analysis we did not attempt to distinguish between these categories, even though our best judgment about the target is indicated on the individual pair maps in the Supplementary material.

The use of acute slices for the functional measurements is another caveat to consider. While we took measures to ensure the health of the recorded neurons, it is possible that some of their axonal or dendritic arbor was damaged or severed during the preparation of live slices. These potential disturbances likely have a minimal effect on the local shortrange neuronal connections that are the focus of this study. On the other hand, the effect of the room temperature conditions during recording are expected to significantly influence the physiological performance of neurons. Thus, at room temperature, conduction velocity is decreased in half (Paintal 1965) or even to one third (Hu and Jonas 2014), and failure rates are higher (Hardingham and Larkman 1998). Therefore the calculated conduction velocity in the present study (0.04 to 0.17 m/s when assuming synaptic delay of 0.25 ms) is likely an underestimate of the actual conduction velocities along PV+ basket cell axons at physiological temperatures, and thus are likely to be at least twice as high as what we observed at room temperature, i.e. in the range of at least 0.1 to 0.35 m/s, or more, bringing it closer to previous measurements. Previous near-physiological temperature experiments estimated the conduction velocity of PV+ basket cell axons in somatosensory cortex of mice to be 0.6 m/s (Casale et al. 2015), and 0.46 m/s in mouse prefrontal cortex (Li et al. 2014). Regarding failure rates, the observed low values at room temperature in our study are expected to be even lower and likely close to 0 at physiological temperature.

One interesting side note from this analysis of synaptic failure rate in pairs of cells making a known number of contacts with each other, is that it gives us an opportunity to calculate an estimate of the failure rate, and accordingly, the release probability (1 – failure rate) for a single inhibitory synapse. Six of our recorded pairs with 8 or fewer synapses between them showed a measureable failure rate. Of course, a ‘failure’ in our recordings is only recorded in a trial where all synapses in the connection fail. If we apply the assumption that all synapses within the pair have the same individual failure rate, the average release probability of a PV+ inhibitory synapse based on our data is 0.28 with a range of 0.09 to 0.43 (see Methods for calculation). Considering that our recordings were done at room temperature this may be an underestimate of the actual release probabilities in physiological conditions (Hardingham and Larkman 1998; Volgushev et al. 2004; but see Pyott and Rosemund 2002). Release probabilities previously estimated for other neocortical synapses vary widely and were shown to depend on the postsynaptic cell type. For example, release probabilities of synapses of layer 2/3 pyramidal neurons vary from 0.13 onto bitufted interneurons, to 0.46 on other pyramidal neurons and 0.64 on multipolar interneurons (Koester and Johnson 2005). The estimated release probabilities of PV+ interneurons in the current study (0.28) are on the lower end of the pyramidal neuron synapses; however, PV+ interneurons make substantially more synapses per connection compared to pyramidal neurons which make 3-5 synapses on average (Deuchars et al. 1994; Feldmeyer et al. 2006), thus increasing the reliability of the connection. There are potentially a number of advantages to having many synapses with low release probabilities in a connection, for example to ensure flexibility and potential for plasticity of the connection, as well as to enable reliable transmission during prolonged high-frequency stimulation (Branco and Staras 2009).

Our study raises an interesting question. If the presence of myelin along the axon correlates with the axonal conduction velocity, then why are the different axonal paths leading to the same postsynaptic target so variable in their myelin coverage? There is no significant correlation between the axonal path length and the extent of its myelin, therefore this will result in a staggered arrival of the input along the different axonal paths within the same connection. This seems contradictory to the known temporal precision of PV+ basket cells neurotransmission. It is plausible that some of the other factors known to influence conduction velocity, such as changes in axonal thickness, length of internodes and nodes of Ranvier length (Debanne et al. 2011; Arancibia-Cárcamo et al. 2017), are also at play, resulting overall in a more unified arrival of the inputs from an individual PV+ basket cell to the postsynaptic target neuron. Indeed, in our electrophysiology recordings, we see no obvious evidence of asynchronous arrive such as ‘notches’ on the rising phase of the IPSC, even in connections with multiple axon paths between the presynaptic PV+ neuron and it’s postsynaptic target. On the other hand, it is tempting to speculate that differences in the conduction velocities of different axonal paths connecting two neurons enable a more nuanced communication, and provide mechanisms through which the inhibitory influence of the PV+ basket cell on its postsynaptic target can be adjusted depending, for example, on the activity of the postsynaptic cell. It is indeed known that PV+ basket connection strength can vary depending on the target neuron activity level (Xue et al. 2014), as well as its outputs (Lee et al. 2014), or whether the connection is reciprocal (Yoshimura and Callaway 2005). Furthermore, PV+ basket cells exhibit an additional level of specificity by placing a portion of their output synapses on more distal dendrites or dendritic spines and enabling local inhibition of excitatory synaptic input to a dendritic branch or a single spine (Somogyi et al. 1983; Kubota et al. 2015). Because different dendrites and dendritic domains of pyramidal neurons vary in their inputs and integration properties (Spruston 2008), PV+ basket cells are thus positioned to control activity levels at vastly different scales – from wide neocortical networks down to specific inputs to individual neurons. Variations in conduction velocity along different axonal paths, and even branch failures under specific conditions, may be one mechanism through which PV+ basket cells accomplish varying levels and patterns of postsynaptic neuron inhibition. There is indeed evidence that axonal branches of the same PV+ basket neuron can be functionally very different, for example by having very different orientation preferences as shown in cat visual cortex (Kisvárday et al. 2002).

In summary, we show here that the presence of myelin along the axons of PV+ interneurons significantly correlates with conduction velocity, underscoring the role of myelin in ensuring fast neurotransmission in short-axon neurons like the PV+ basket cell in neocortex, in addition to its well-known role for long-range projection neurons. Axonal myelination appears to be part of a coordinated array of PV+ interneuron specializations that also include, as shown here, redundancy of its individual connections that are formed through multiple axonal paths targeting the postsynaptic neuron onto multiple domains, as well as through convergence of several axonal paths onto the same postsynaptic domain. One of the main roles that has been ascribed to cortical interneurons has been the influence and control over rhythmic activity in cortical circuits, in particular gamma rhythms (Cardin et al. 2009; Buzsáki and Wang 2012; Cardin 2018; Chariker et al. 2018). Inhibitory basket neurons have electrophysiological properties that tend to entrain their activity at gamma rhythms, and the time course of GABAA mediated IPSPs are optimal for supporting rhythms in the gamma range. The factors generating these rhythms are complex, being influenced strongly by excitatory/inhibitory ratio in cortical circuits (Yizhar et al. 2011; Cardin 2018), but are directly controlled by the activity of PV+ interneurons (Sohal et al. 2009; Orekhova et al. 2015; Antonoudiou et al. 2020), occur in a frequency range of 30-80 Hz (Cardin 2018), and thus have a periodicity in the range of ~10-30 ms. Our findings suggest that myelination of PV+ interneurons can decrease the latency of the synaptic response they generate on the order of a millisecond. This would represent a potential phase shift in a gamma-type rhythm of 3-8%, an amount that is certainly large enough to influence neuronal circuit functions that rest upon these rhythmic activities. Potential phase shifts of this magnitude may well have relevance for neural circuit performance and behavior (Cardin et al. 2009; Sohal et al. 2009; Yizhar et al. 2011; Orekhova et al. 2015; Kim et al. 2016; Cardin 2018). If later research were to reveal that the extent of myelination on PV+ interneuron axons is plastic in a reasonably short time frame, this could also represent an important novel mechanism for controlling the interaction between these interneurons and the neuronal functions they subserve, including the behaviorally relevant regulation of cortical rhythms.

## Supporting information

Supplement 1: axonograms

Supplement 2: dendrograms

## Acknowledgements

We thank Sonja Winter from the Department of Psychological Sciences at UC Merced for helpful discussion.

