## Supplement 1: axonograms for "Conduction velocity along the local axons of parvalbumin interneurons correlates with the degree of axonal myelination"

### Supplementary material: Axonograms

#### Legend:

*Cell body*

*Axon*

*Myelin*

*Incomplete myelination*

*Untraced*

***AP1*** *Axonal path 1*

*Contact on dendritic spine*

*Contact on dendritic shaft*

*Contact on soma*

*Contact on AIS*

17.4: length in μm, not corrected for shrinkage

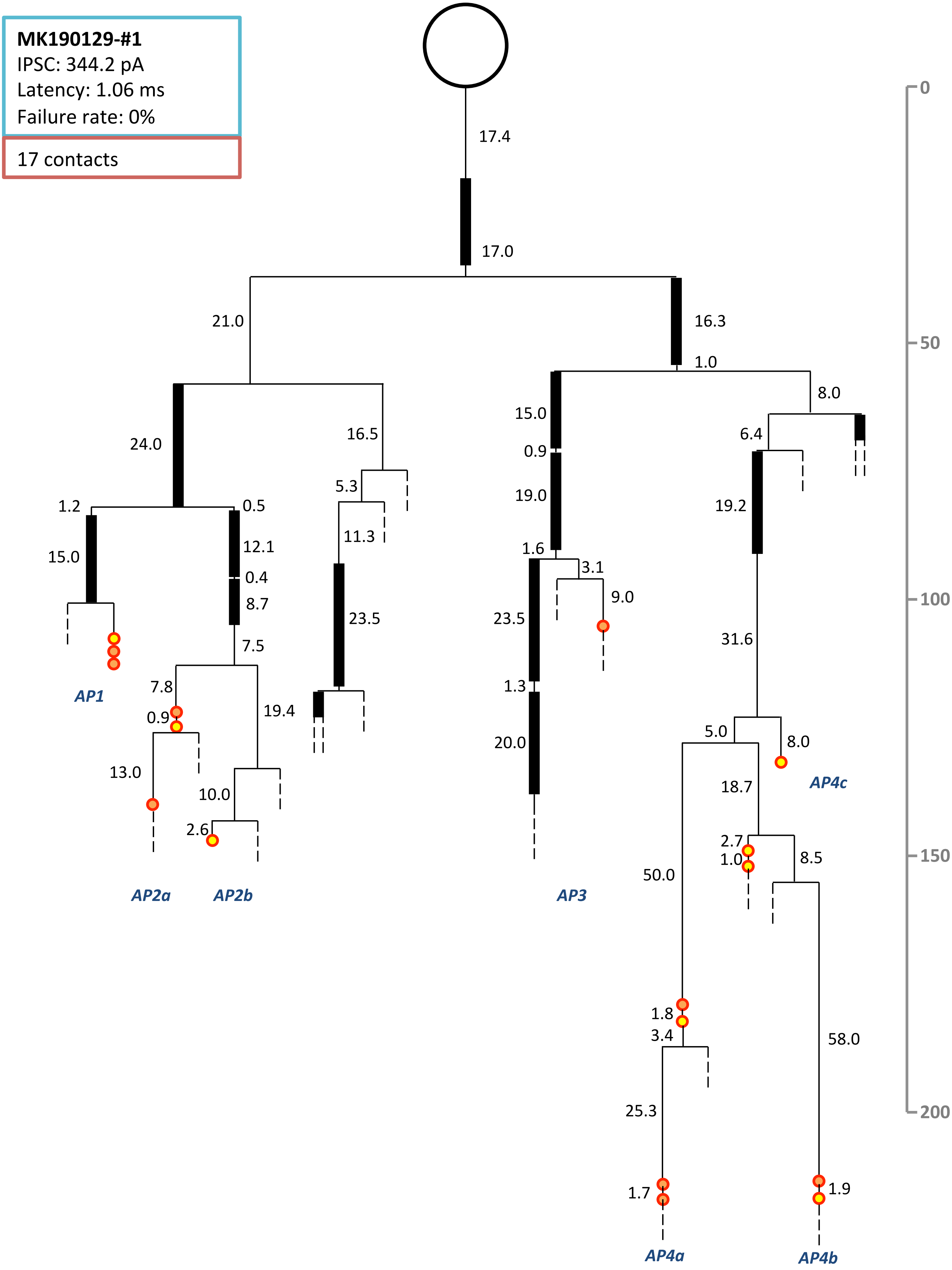
MK190129-#1
IPSC: 344.2 pA
Latency: 1.06 ms
Failure rate: 0%
17 contacts
17.4
17.0
21.0
16.3
1.0
8.0
24.0
16.5
15.0
0.9
6.4
5.3
19.0
19.2
1.2
0.5
12.1
11.3
1.6
3.1
15.0
0.4
9.0
8.7
23.5
23.5
7.5
31.6
AP1
7.8
19.4
1.3
0.9
5.0
8.0
20.0
13.0
AP4c
10.0
18.7
2.6
2.7
1.0
8.5
AP2a
AP2b
AP3
50.0
1.8
3.4
58.0
25.3
1.7
1.9
AP4a
AP4b
0
50
100
150
200

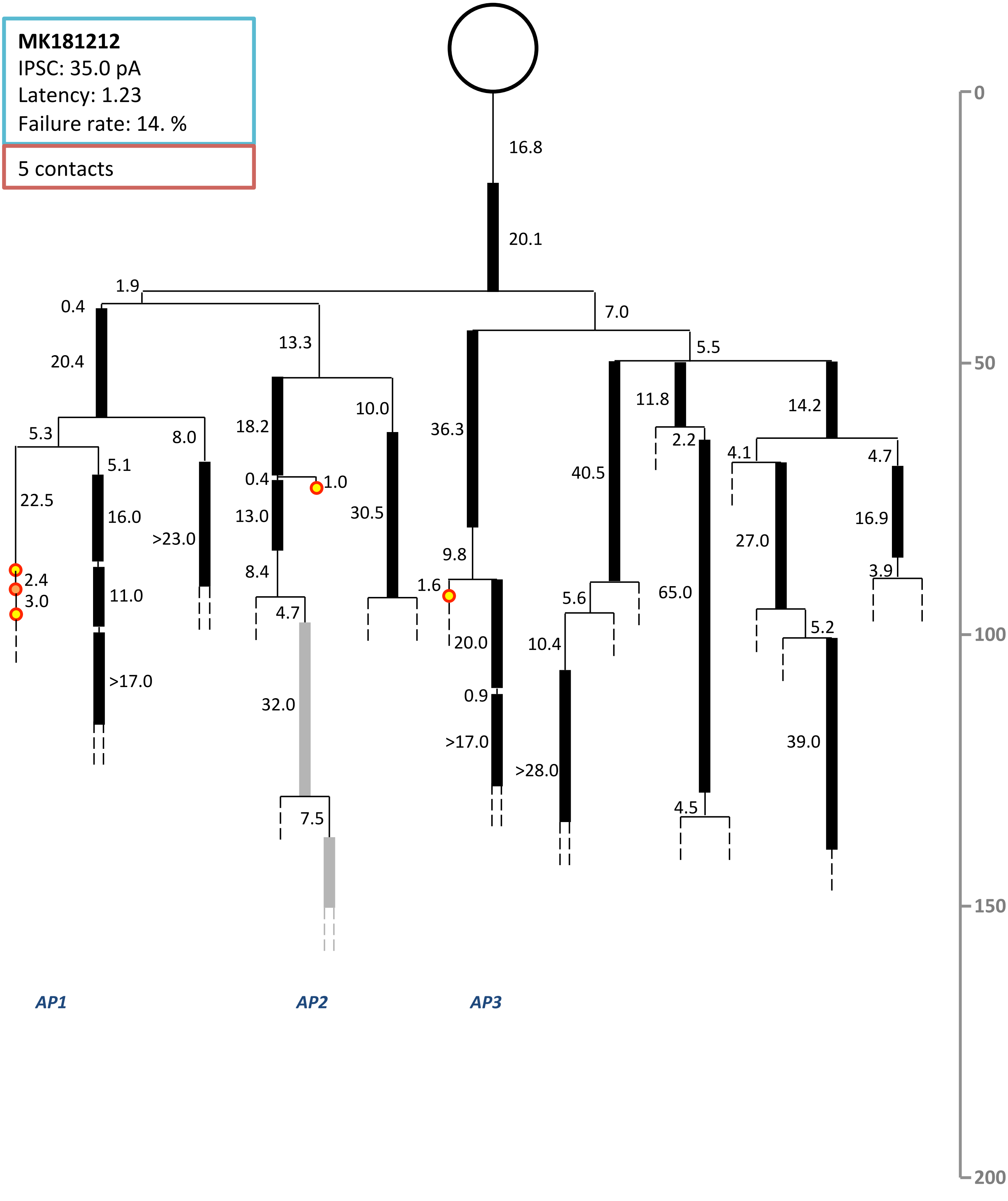
MK181212
IPSC: 35.0 pA
Latency: 1.23
Failure rate: 14. %
5 contacts
16.8
20.1
1.9
0.4
20.4
7.0
13.3
5.5
5.3
8.0
5.1
22.5
16.0
>23.0
2.4
3.0
11.0
>17.0
18.2
10.0
0.4
1.0
13.0
30.5
8.4
4.7
32.0
7.5
36.3
9.8
1.6
20.0
0.9
>17.0
40.5
5.6
10.4
>28.0
11.8
2.2
65.0
4.5
14.2
4.1
4.7
27.0
16.9
3.9
5.2
39.0
AP1
AP2
AP3
0
50
100
150
200

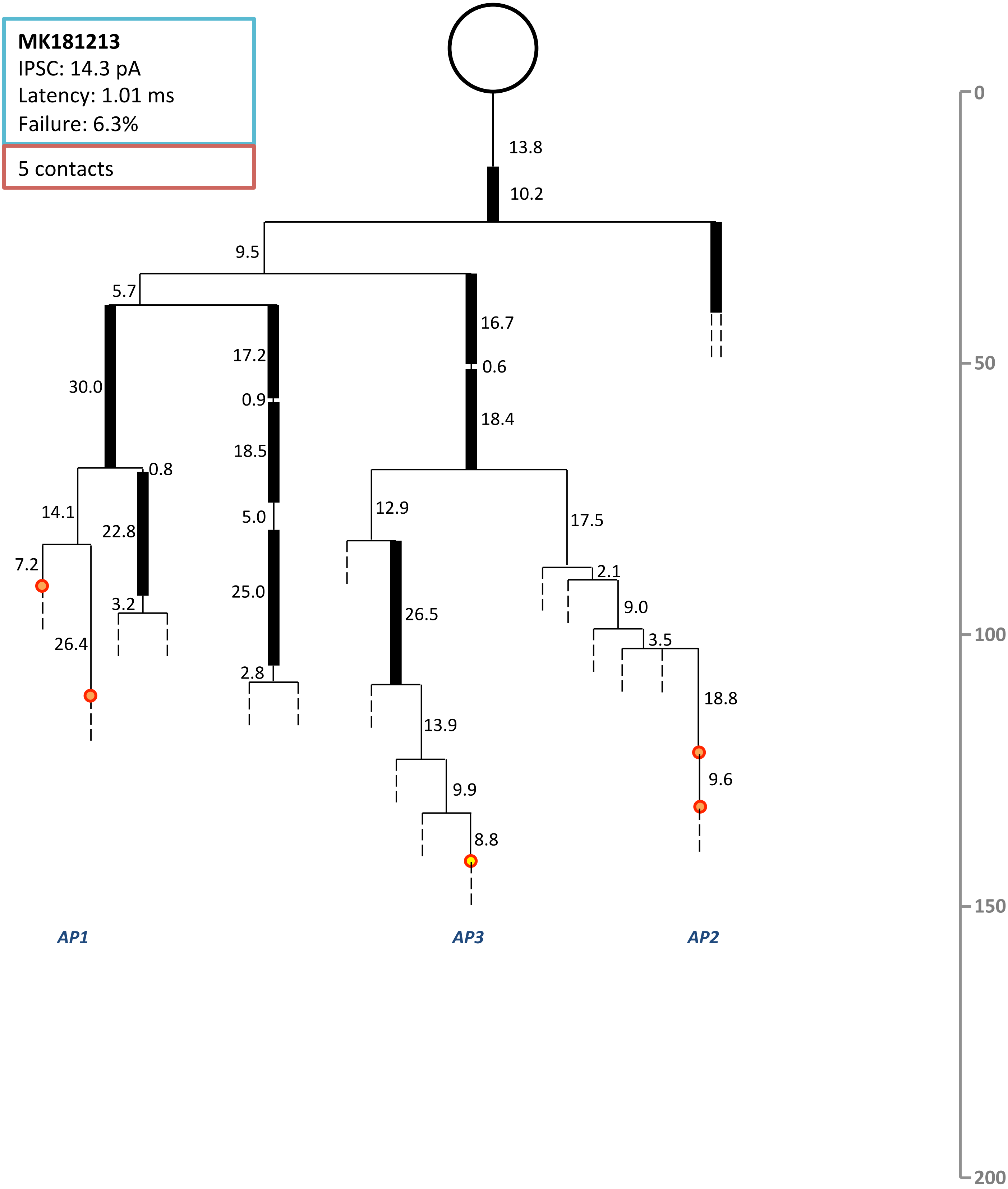

MK181213
IPSC: 14.3 pA
Latency: 1.01 ms
Failure: 6.3%
5 contacts
13.8
10.2
9.5
5.7
16.7
0.6
18.4
30.0
17.2
0.9
18.5
0.8
14.1
22.8
5.0
12.9
17.5
7.2
3.2
25.0
26.5
2.1
9.0
3.5
26.4
2.8
18.8
13.9
9.9
9.6
8.8
AP1
AP3
AP2
0
50
100
150
200

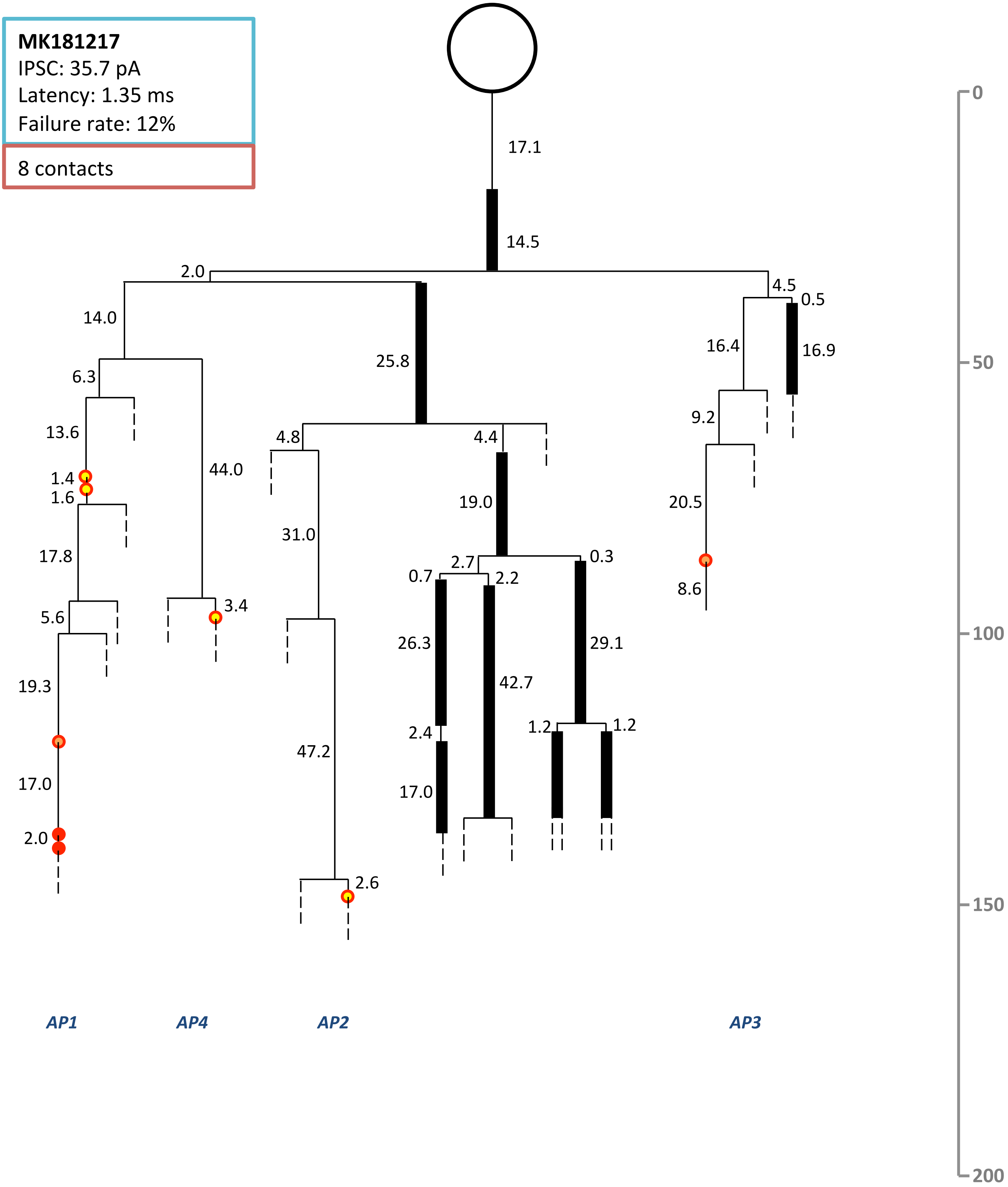

MK181217
IPSC: 35.7 pA
Latency: 1.35 ms
Failure rate: 12%
8 contacts
17.1
14.5
2.0
4.5
0.5
14.0
25.8
16.4
16.9
6.3
9.2
13.6
4.8
4.4
44.0
1.4
1.6
19.0
20.5
31.0
17.8
2.7
0.3
0.7
2.2
8.6
5.6
3.4
26.3
29.1
42.7
19.3
2.4
1.2
1.2
47.2
17.0
17.0
2.0
2.6
AP1
AP4
AP2
AP3
0
50
100
150
200

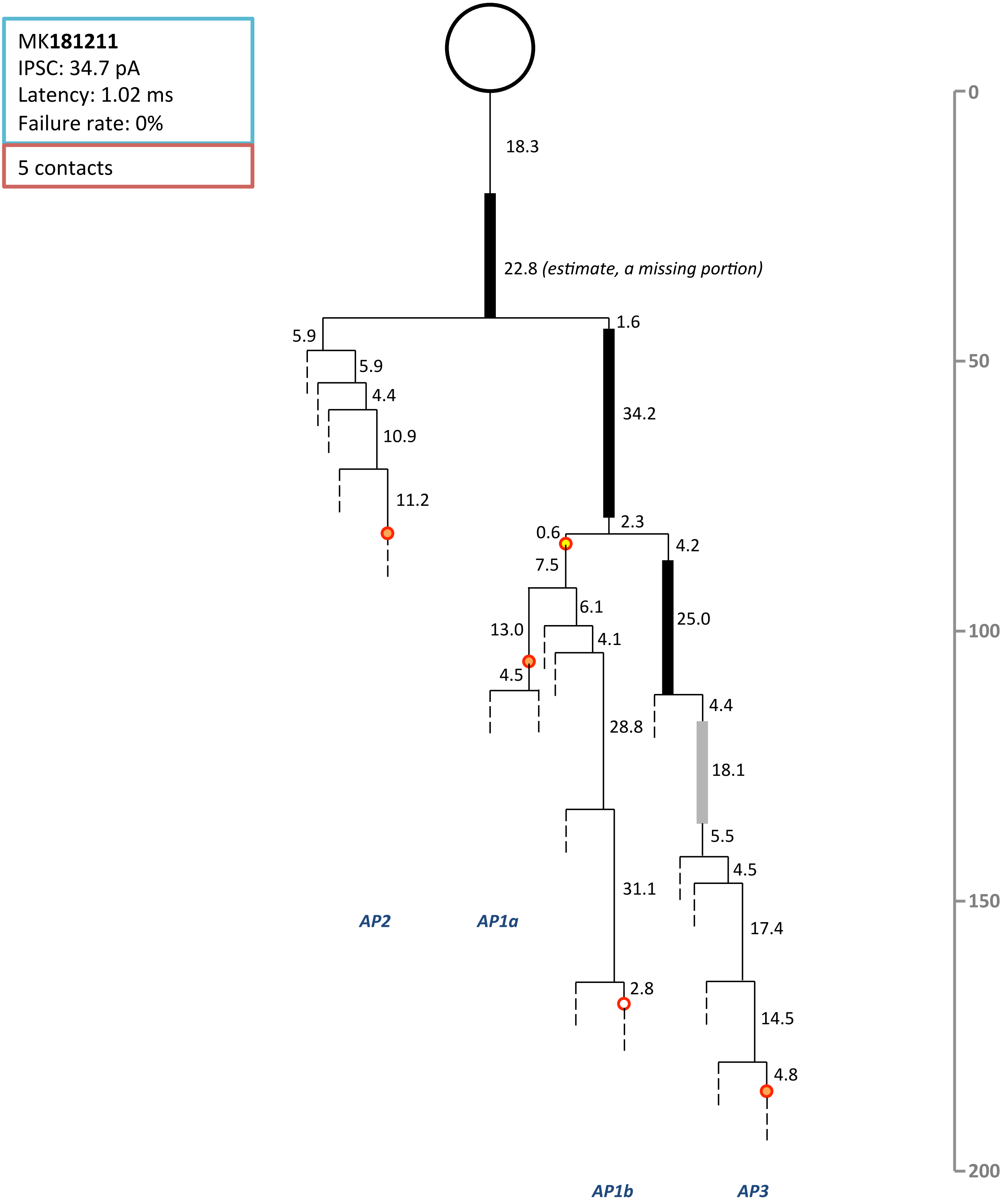

MK181211
IPSC: 34.7 pA
Latency: 1.02 ms
Failure rate: 0%
5 contacts
18.3
22.8 (estimate, a missing portion)
5.9
1.6
5.9
4.4
34.2
10.9
11.2
0.6
2.3
4.2
7.5
6.1
25.0
13.0
4.1
4.5
4.4
28.8
18.1
5.5
4.5
31.1
AP2
AP1a
17.4
2.8
14.5
4.8
AP1b
AP3
0
50
100
150
200

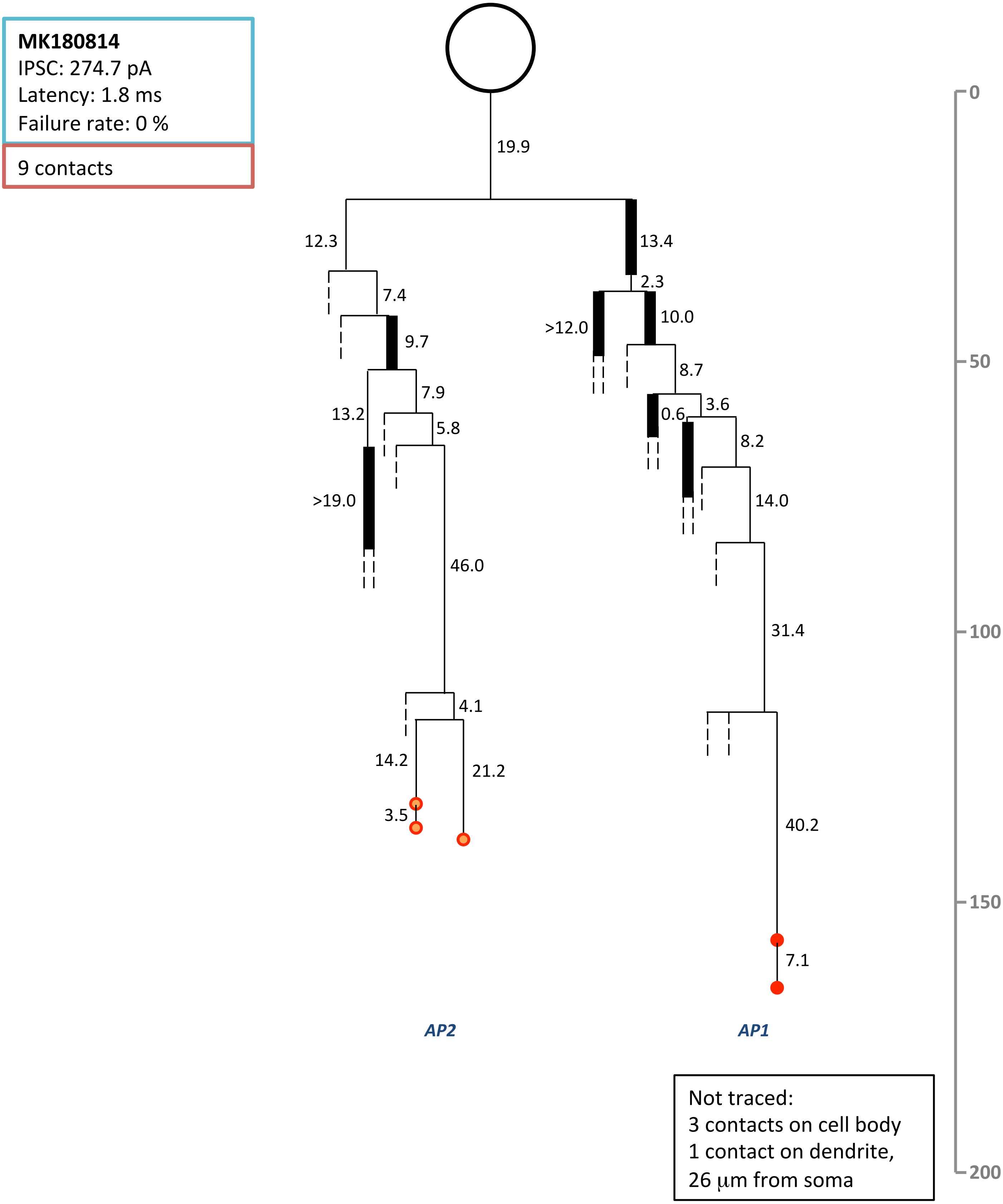
MK180814
IPSC: 274.7 pA
Latency: 1.8 ms
Failure rate: 0 %
9 contacts
19.9
12.3
13.4
7.4
2.3
9.7
>12.0
10.0
7.9
8.7
13.2
5.8
0.6
3.6
8.2
>19.0
14.0
46.0
31.4
4.1
14.2
21.2
3.5
40.2
7.1
AP2
AP1
Not traced:
3 contacts on cell body
1 contact on dendrite,
26 µm from soma
0
50
100
150
200

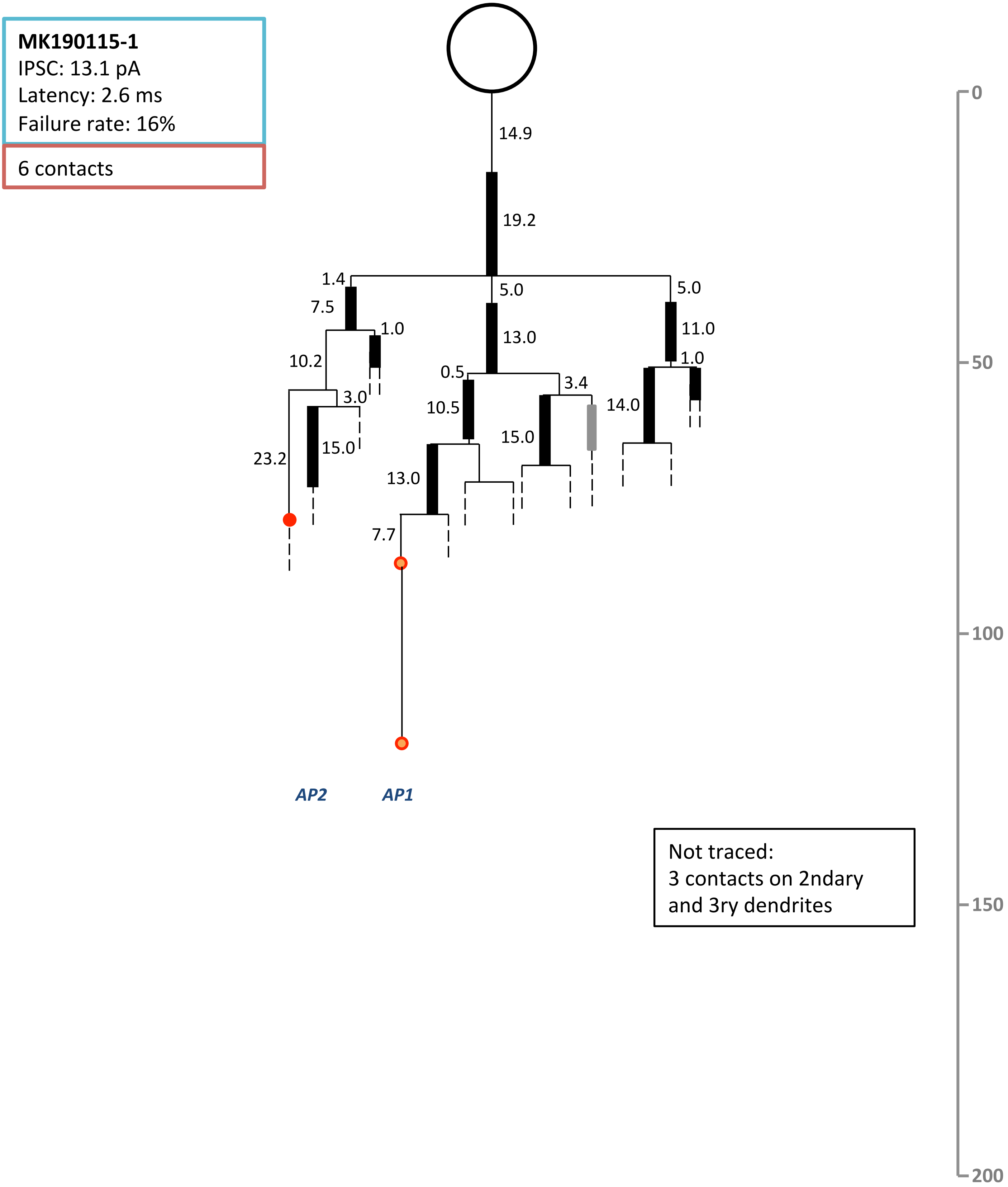

MK190115-1
IPSC: 13.1 pA
Latency: 2.6 ms
Failure rate: 16%
6 contacts
14.9
19.2
1.4
5.0
5.0
7.5
1.0
13.0
11.0
10.2
0.5
1.0
3.0
3.4
10.5
14.0
15.0
15.0
23.2
13.0
7.7
AP2
AP1
Not traced:
3 contacts on 2ndary
and 3ry dendrites
0
50
100
150
200

**MK190110**
IPSC: 33.8 pA
Latency:1.8 ms
Failure rate: 12.5%

5 contacts

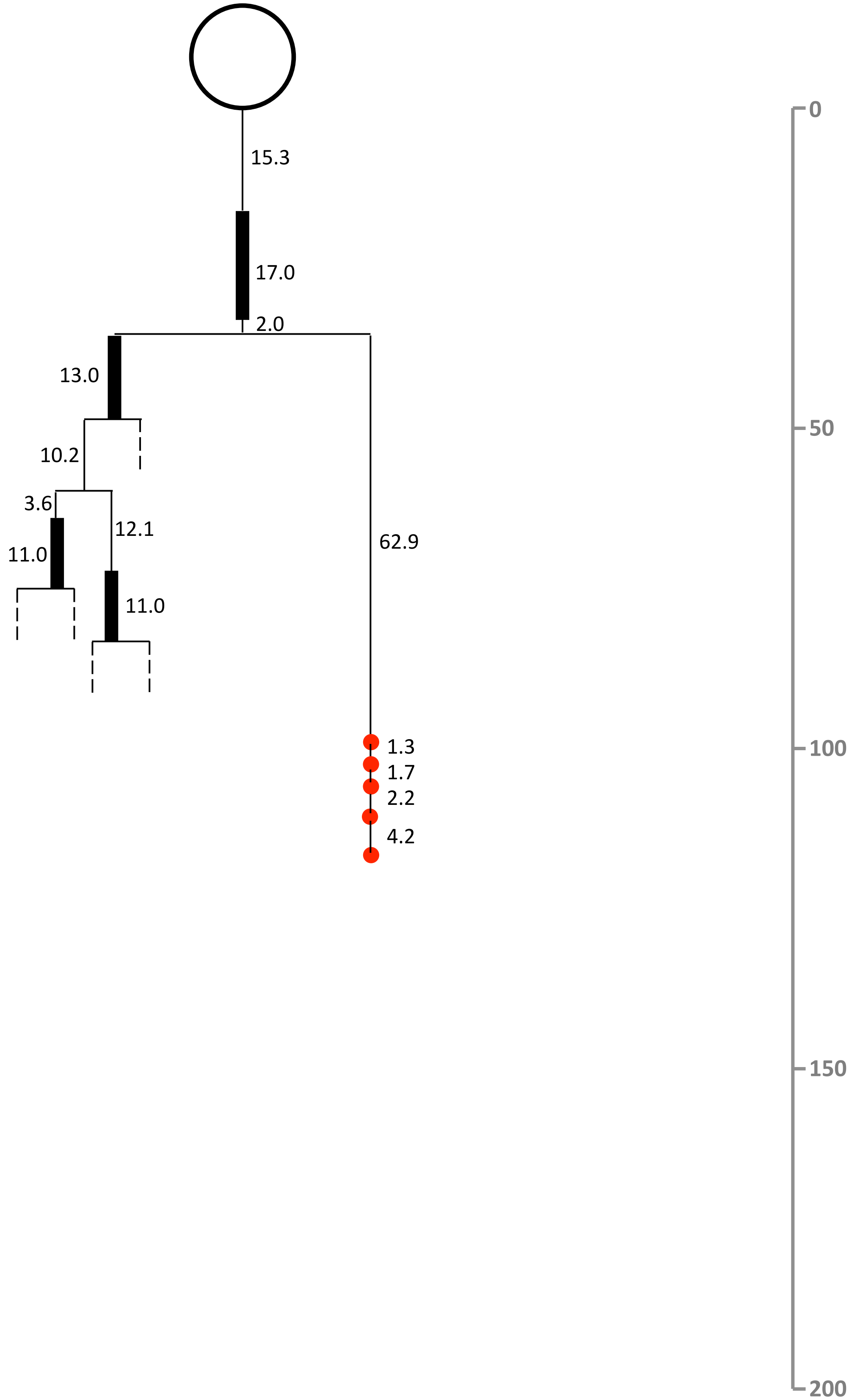

**MK190227**
IPSC: 300.1 pA
Latency: 1.38 ms
Failure rate: 0%

18 contacts

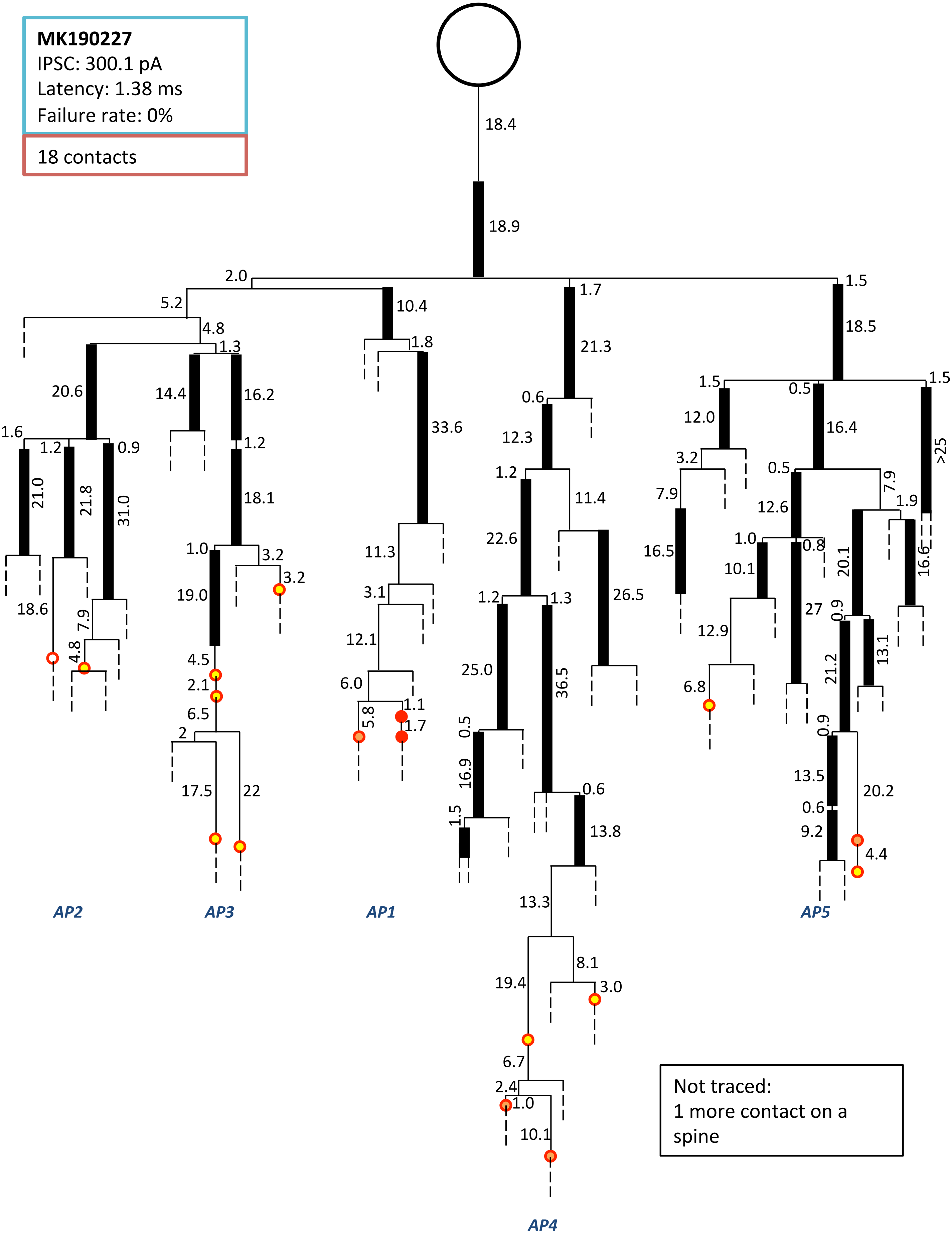


Not traced:
1 more contact on a spine

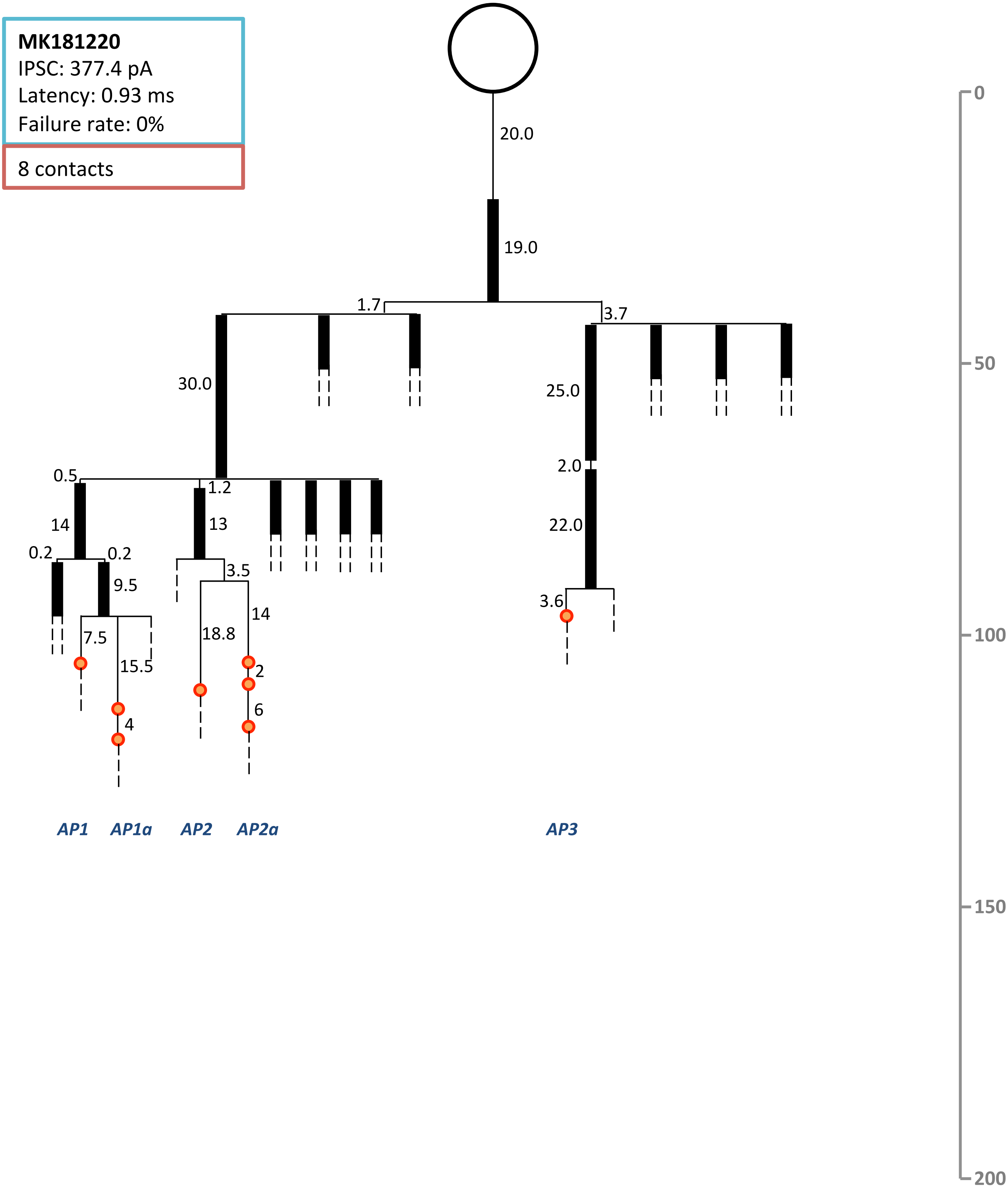
MK181220
IPSC: 377.4 pA
Latency: 0.93 ms
Failure rate: 0%
8 contacts
20.0
19.0
1.7
3.7
30.0
25.0
2.0
22.0
0.5
1.2
14
13
0.2
0.2
9.5
3.5
7.5
15.5
18.8
14
2
6
4
3.6
AP1
AP1a
AP2
AP2a
AP3
0
50
100
150
200

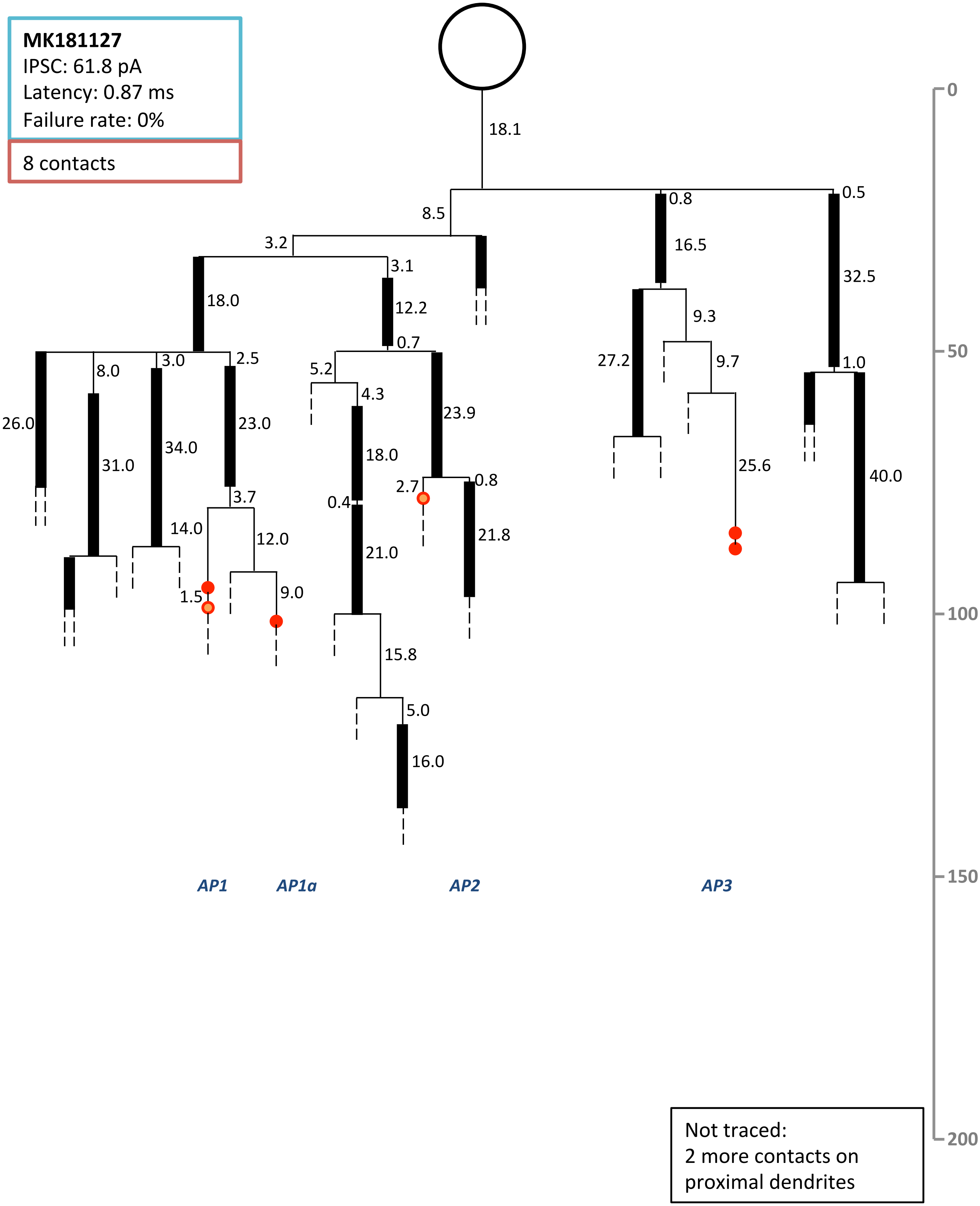
**MK181127**
IPSC: 61.8 pA
Latency: 0.87 ms
Failure rate: 0%
8 contacts
AP1
AP1a
AP2
AP3
0
50
100
150
200
Not traced:
2 more contacts on
proximal dendrites

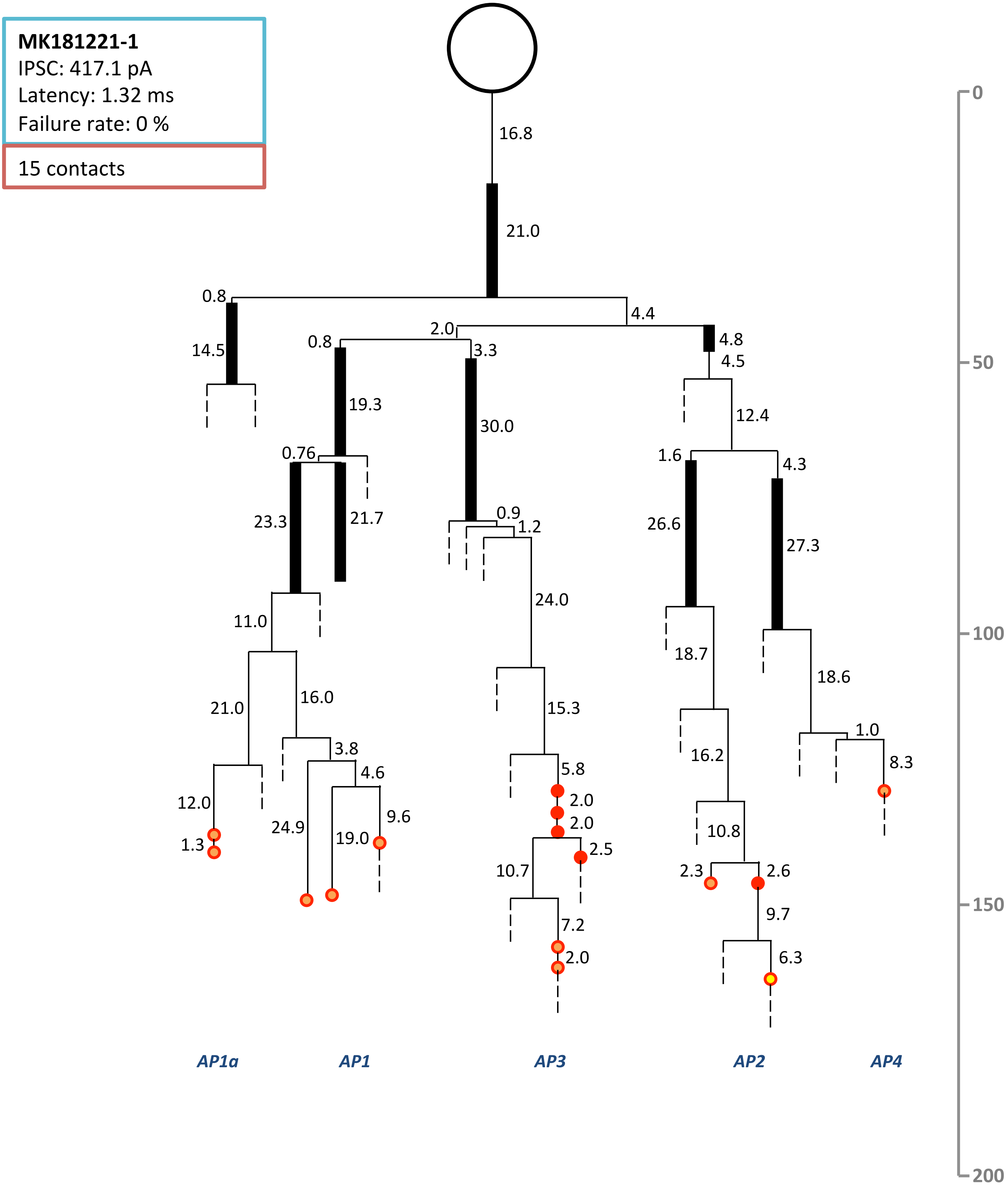

MK181221-1
IPSC: 417.1 pA
Latency: 1.32 ms
Failure rate: 0 %
15 contacts
16.8
21.0
0.8
4.4
14.5
2.0
0.8
3.3
4.8
4.5
19.3
30.0
12.4
0.76
1.6
4.3
23.3
21.7
0.9
1.2
26.6
27.3
24.0
11.0
18.7
18.6
21.0
16.0
15.3
16.2
3.8
1.0
4.6
5.8
8.3
12.0
2.0
2.0
9.6
24.9
19.0
10.8
1.3
2.5
10.7
2.3
2.6
9.7
7.2
2.0
6.3
AP1a
AP1
AP3
AP2
AP4
0
50
100
150
200

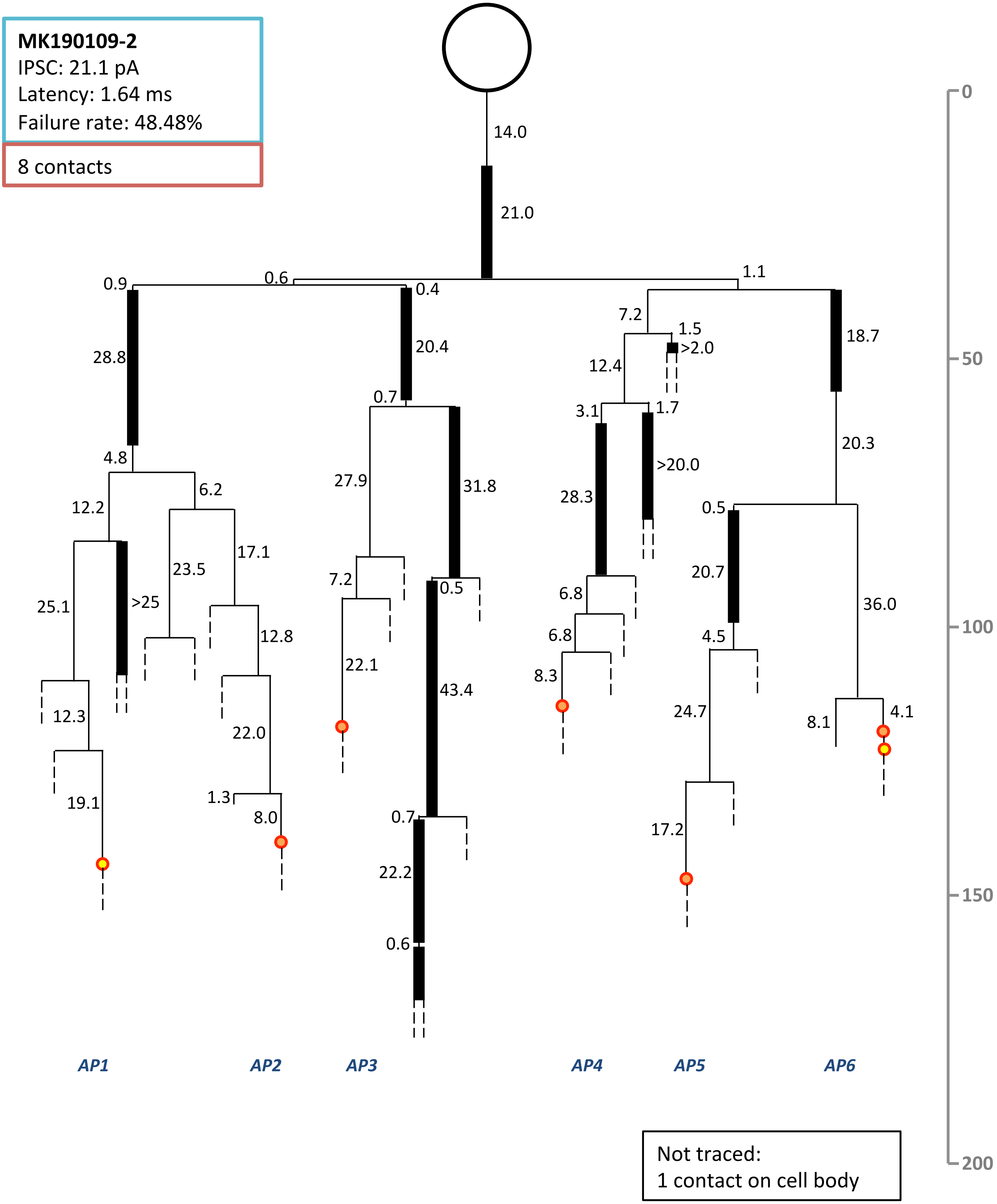
MK190109-2
IPSC: 21.1 pA
Latency: 1.64 ms
Failure rate: 48.48%
8 contacts
14.0
21.0
0.6
1.1
0.9
0.4
28.8
20.4
7.2
1.5
>2.0
18.7
12.4
0.7
3.1
1.7
4.8
6.2
12.2
27.9
31.8
>20.0
28.3
20.3
0.5
17.1
23.5
25.1
>25
7.2
0.5
6.8
20.7
36.0
12.8
6.8
4.5
22.1
43.4
8.3
12.3
24.7
8.1
4.1
22.0
19.1
1.3
8.0
0.7
17.2
22.2
0.6
AP1
AP2
AP3
AP4
AP5
AP6
0
50
100
150
200
Not traced:
1 contact on cell body
