## Supplement 2: dendrograms for "Conduction velocity along the local axons of parvalbumin interneurons correlates with the degree of axonal myelination"

### Supplementary material: Dendrograms

Legend:

*Cell body*

*Primary dendrite*

*Axon initial segment*

*Secondary dendrite*

*Tertiary dendrite*

*Contact on spine*

*Contact on dendr. shaft*

*Contact on soma*

*Contact on AIS*

***AP1*** *Axonal path 1*

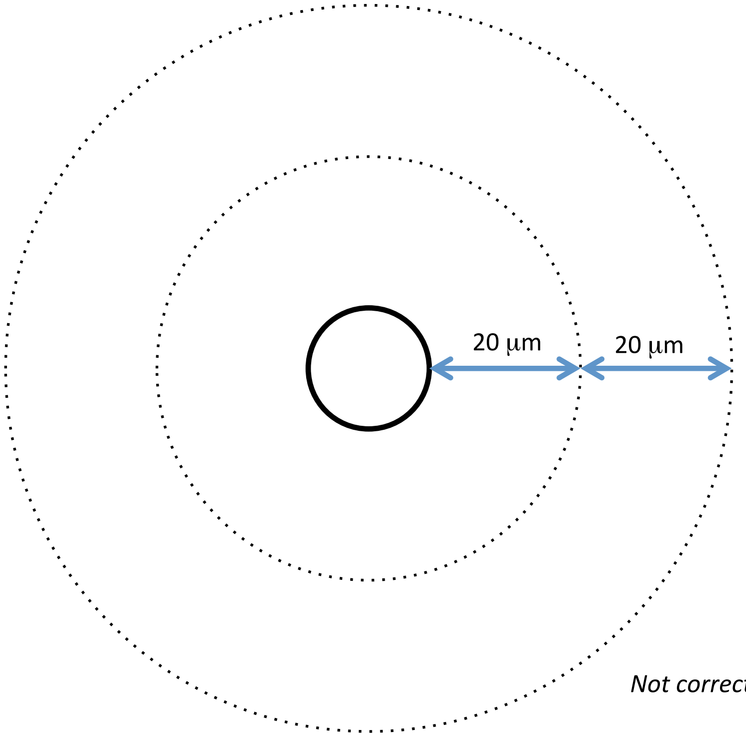


*Not corrected for shrinkage*

**MK190129-#1**
IPSC: 344.2 pA
Latency: 1.06 ms
Failure rate: 0%

17 contacts

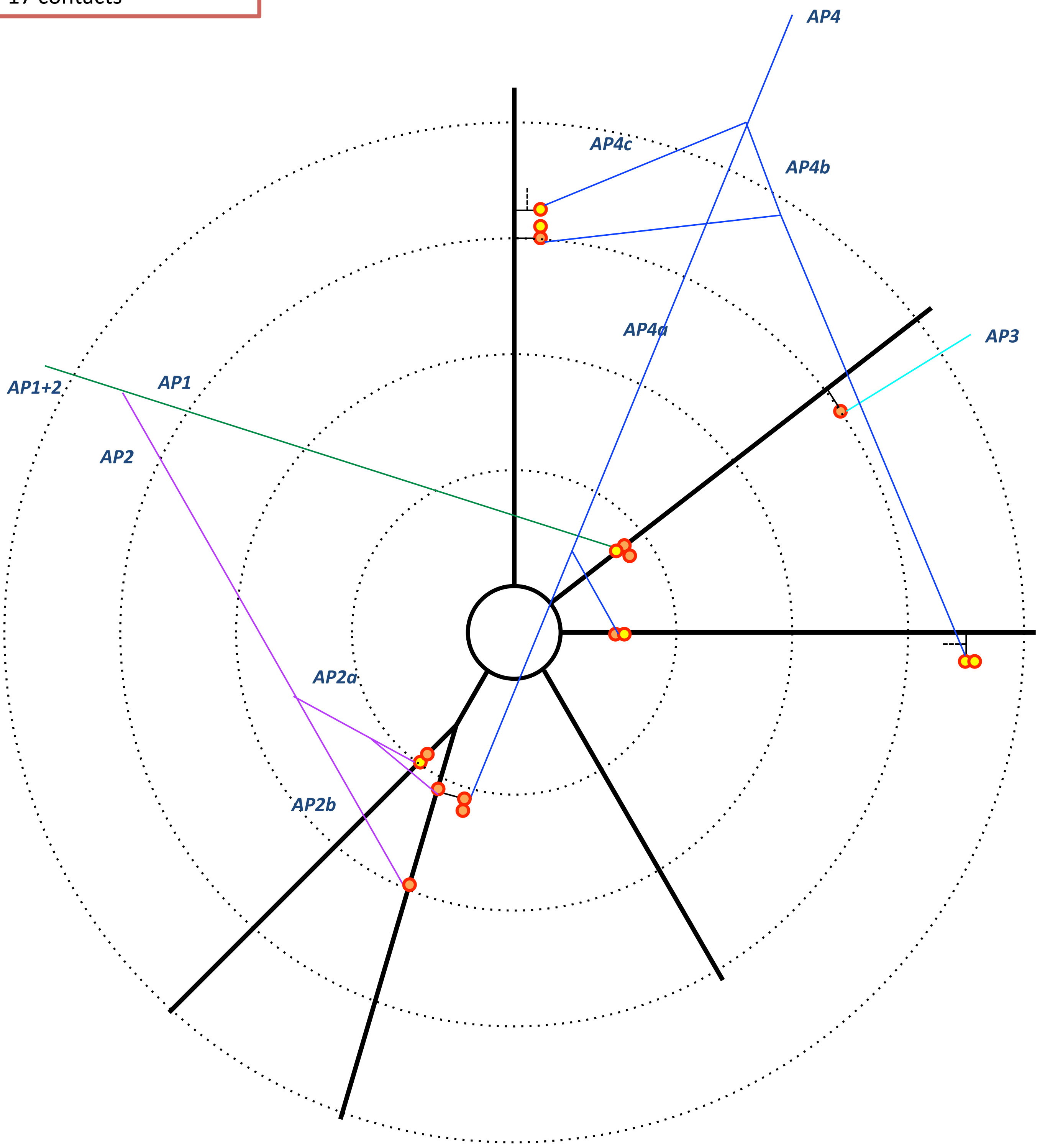

**MK181212**
IPSC: 35.0 pA
Latency: 1.23
Failure rate: 14. %

5 contacts

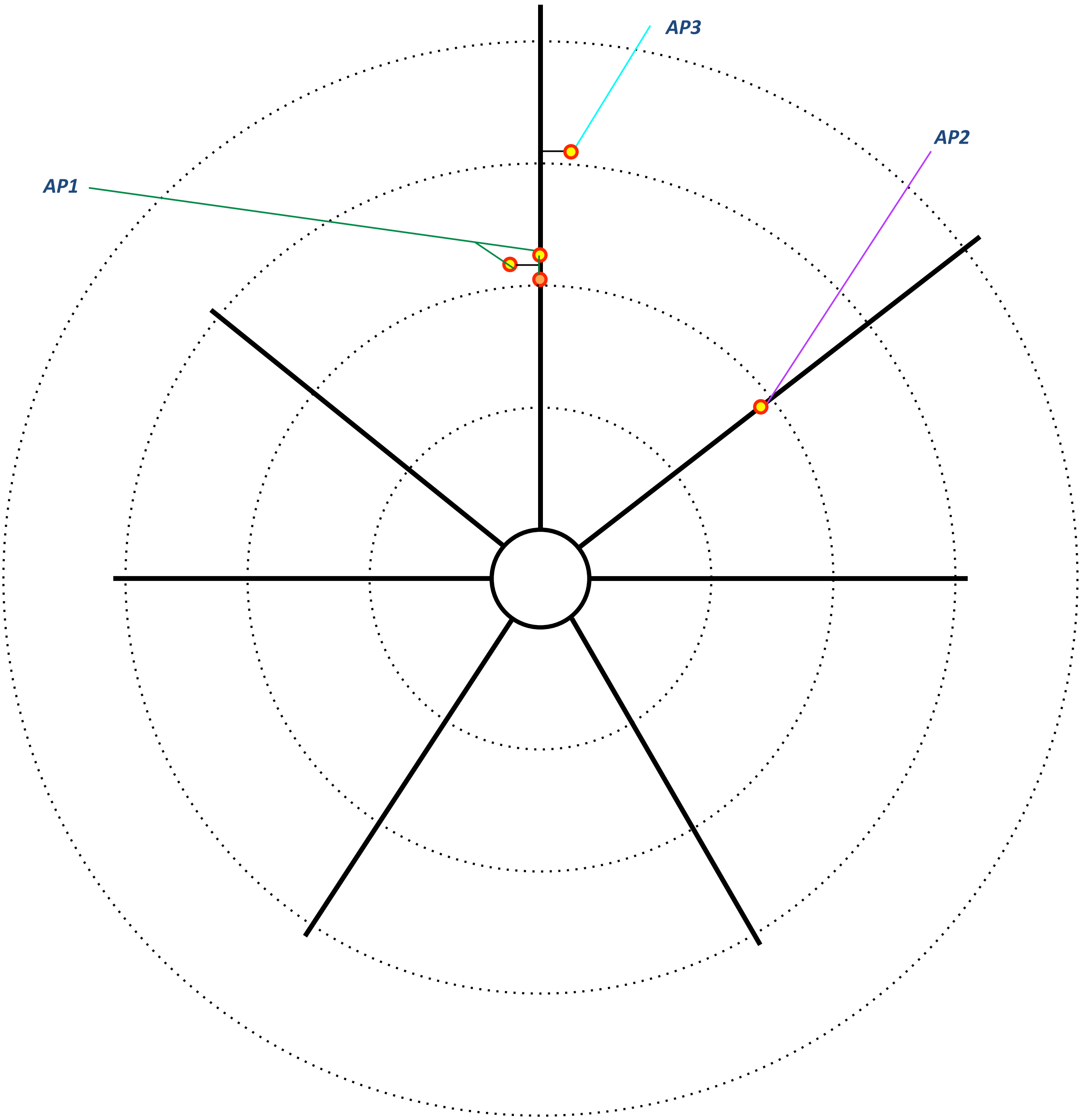

**MK181213**
IPSC: 14.3 pA
Latency: 1.01 ms
Failure: 6.3%

5 contacts

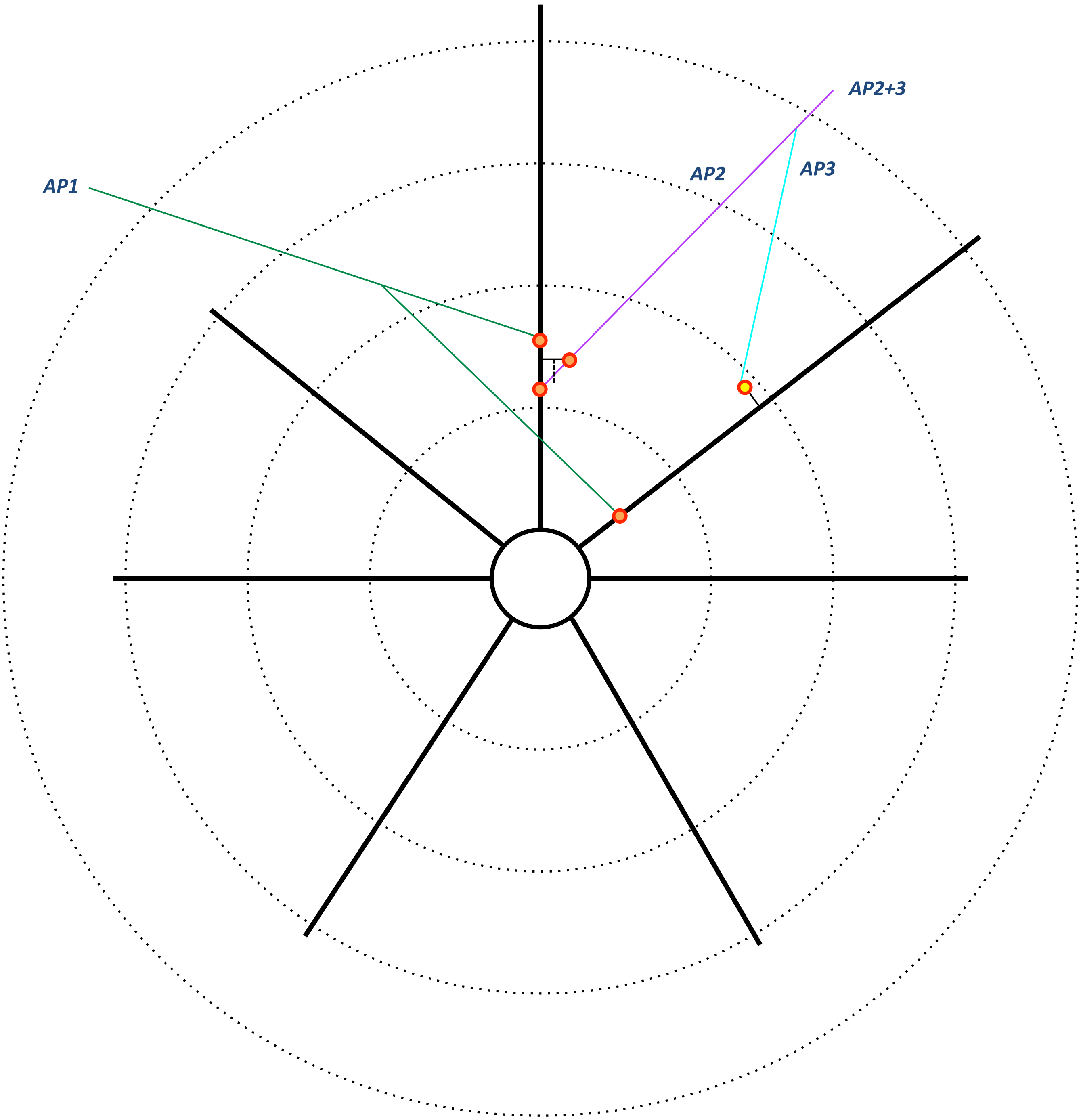

**MK181217**
IPSC: 35.7 pA
Latency: 1.35 ms
Failure rate: 12%

8 contacts

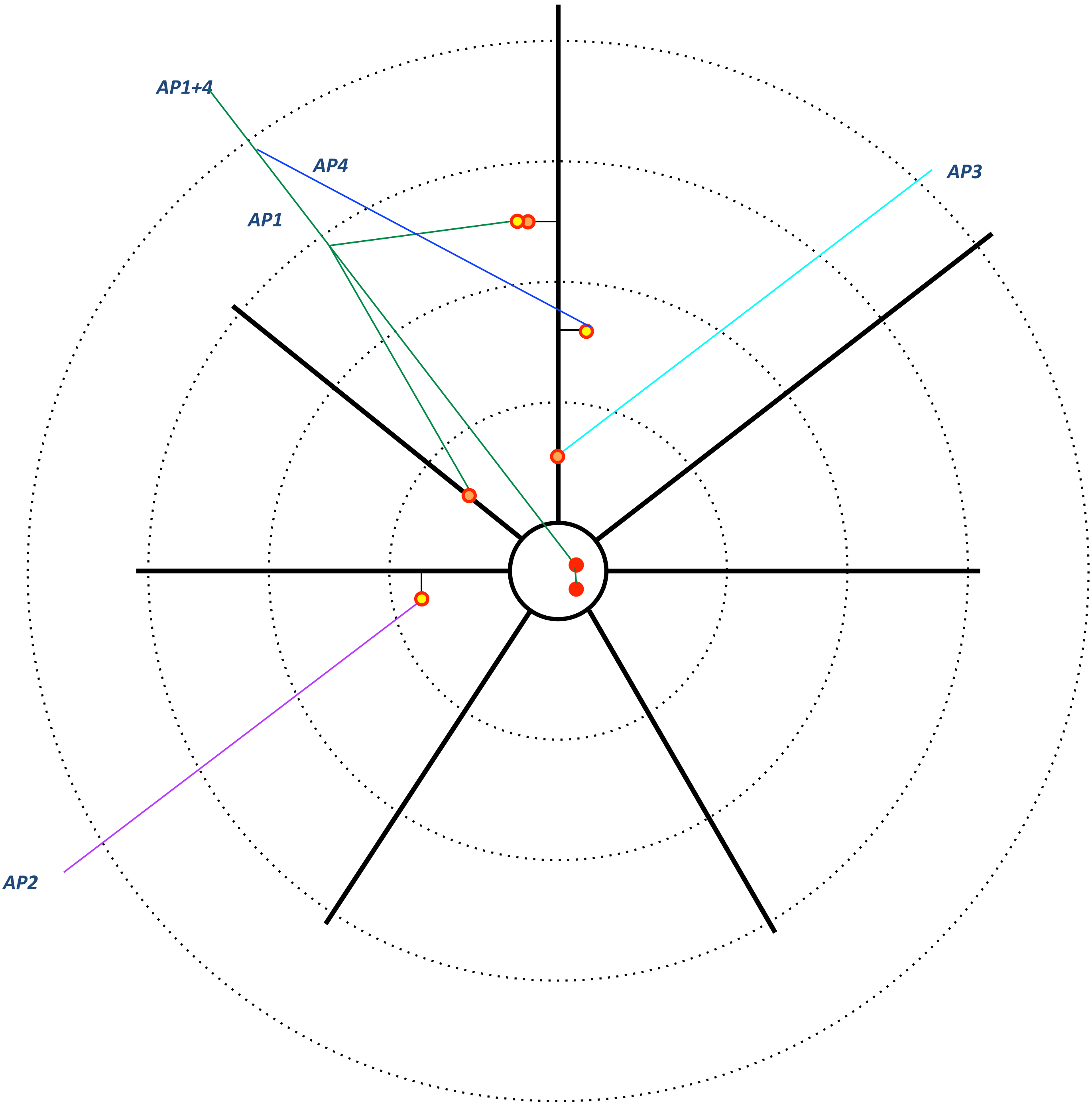

MK**181211**
IPSC: 34.7 pA
Latency: 1.02 ms
Failure rate: 0%

5 contacts

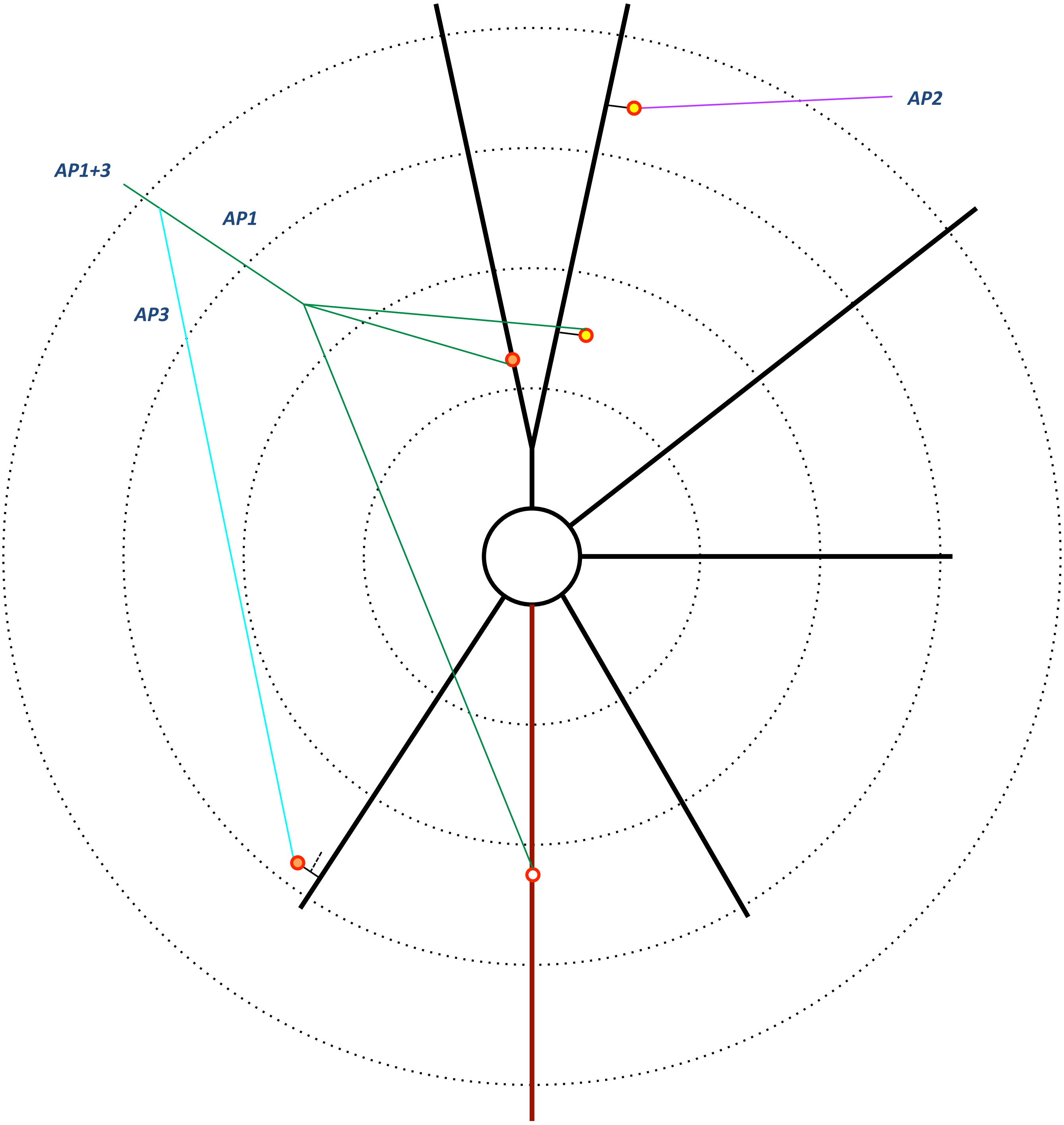

**MK180814**
IPSC: 274.7 pA
Latency: 1.8 ms
Failure rate: 0 %

9 contacts

Interneuron

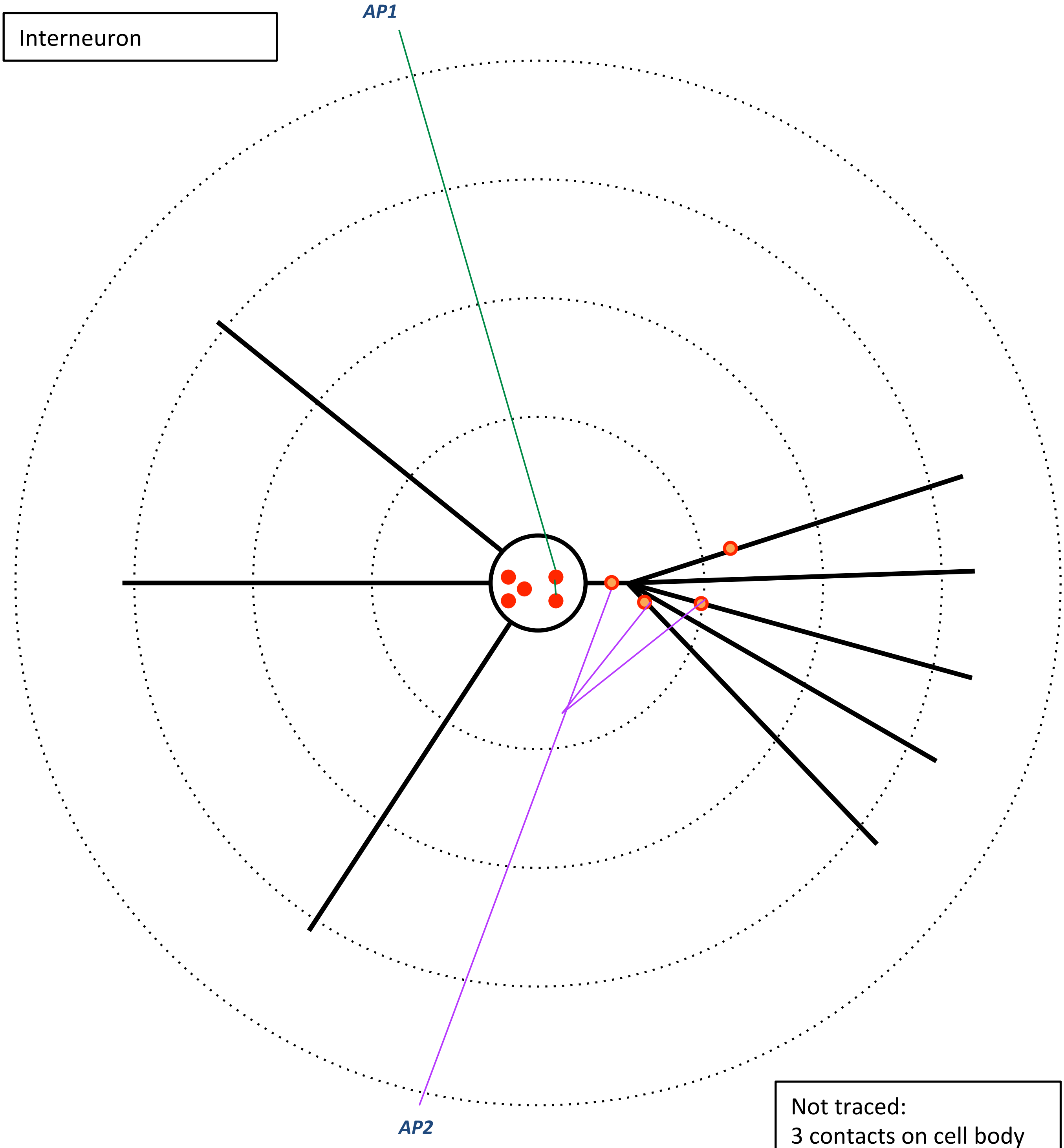


Not traced:
3 contacts on cell body
1 contact on dendrite,
26 μm from soma

**MK190115-1**
IPSC: 13.1 pA
Latency: 2.6 ms
Failure rate: 16%

6 contacts

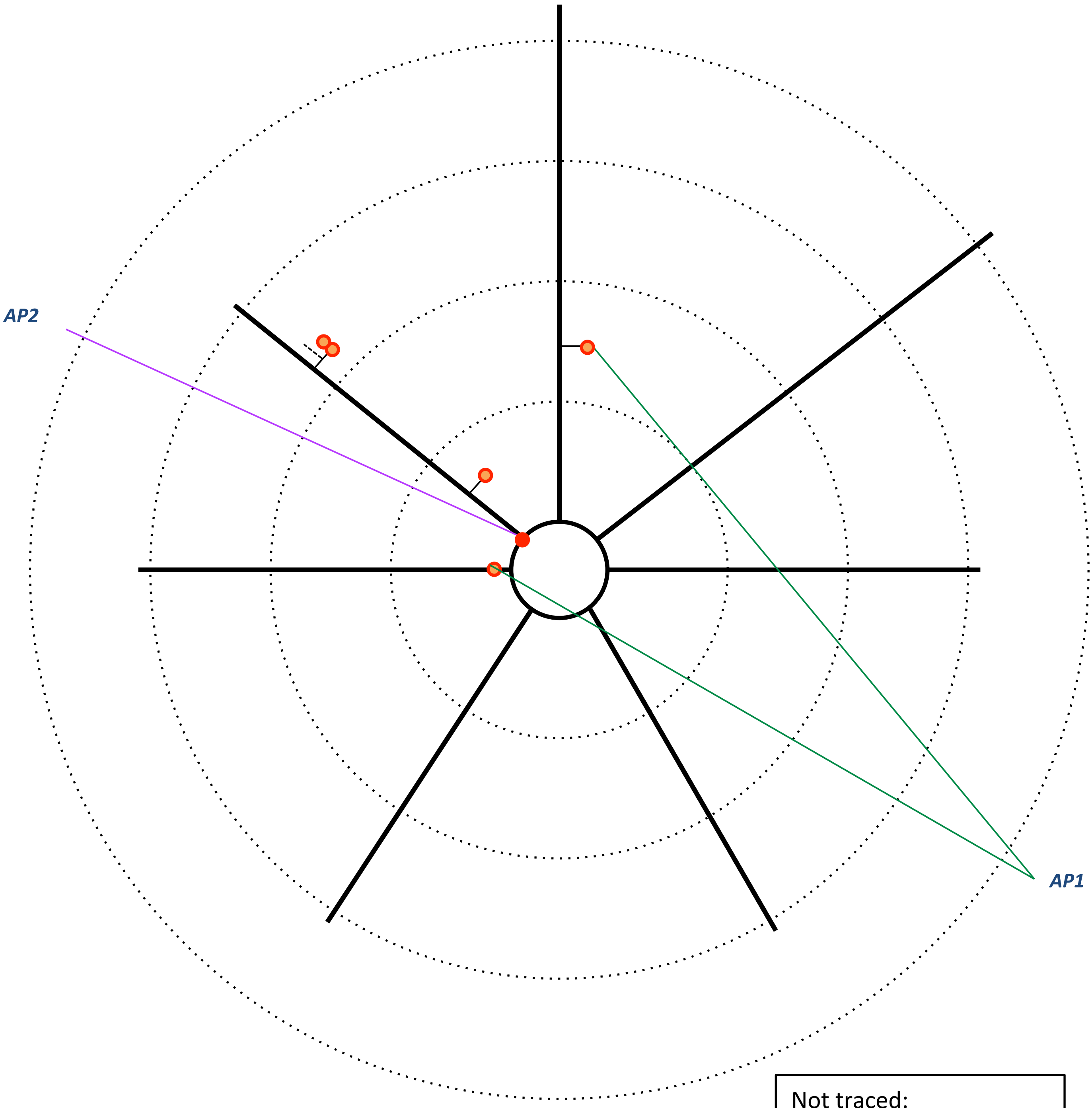


Not traced:
3 contacts on 2ndary and 3ry dendrites

**MK190110**
IPSC: 33.8 pA
Latency:1.8 ms
Failure rate: 12.5%

5 contacts

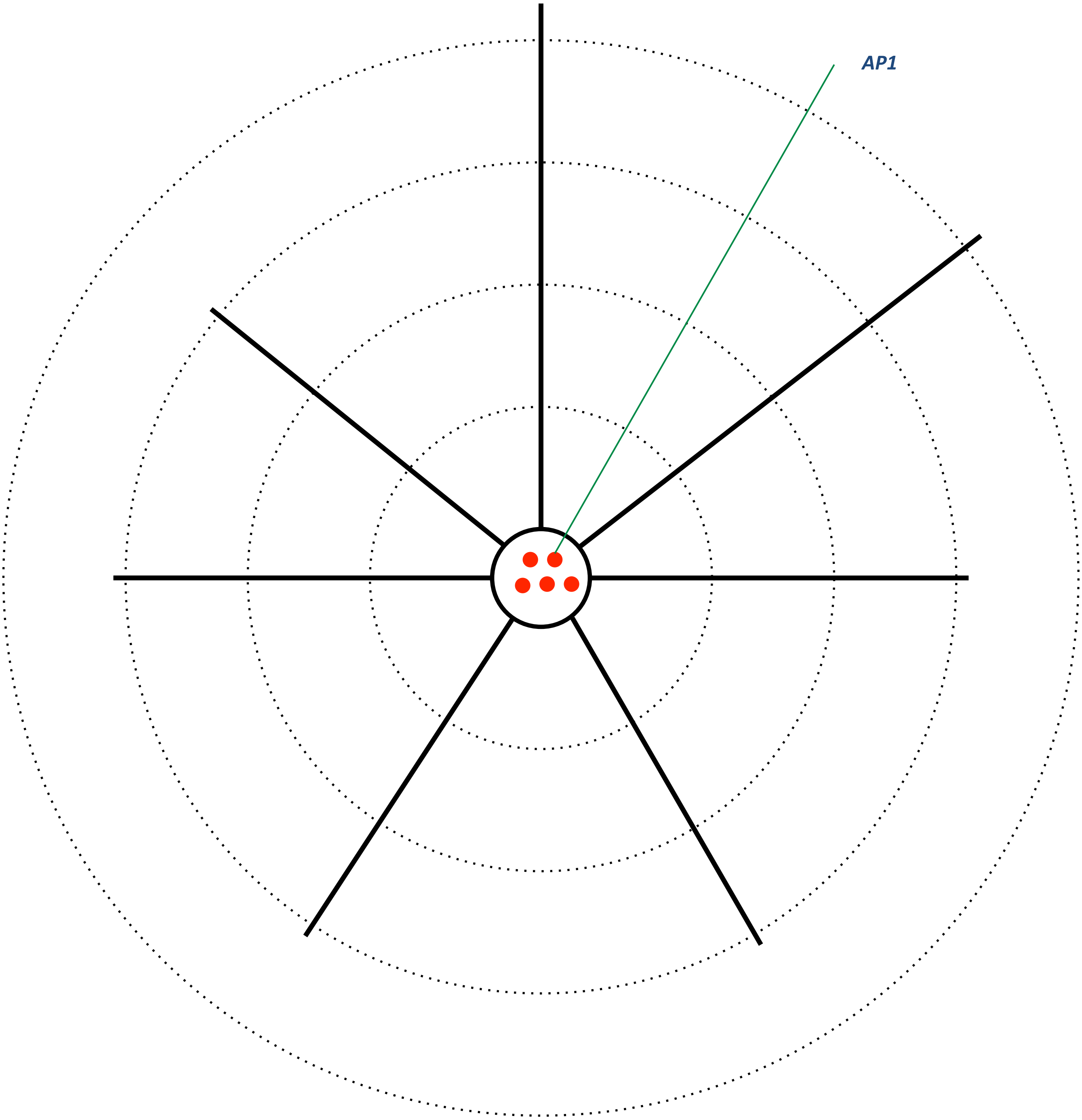

**MK190227**
IPSC: 300.1 pA
Latency: 1.38 ms
Failure rate: 0%

18 contacts

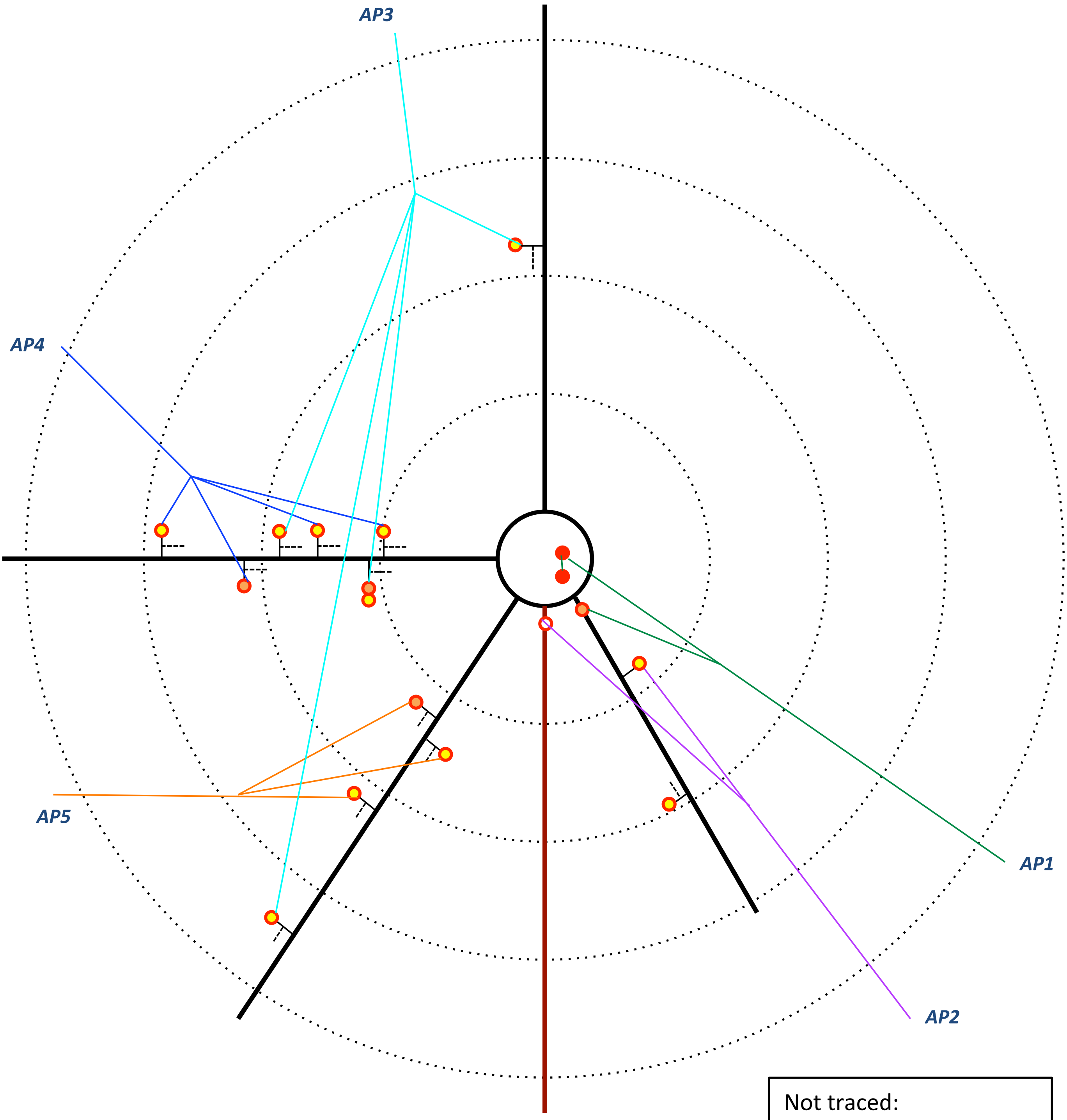


Not traced:
1 more contact on a spine

**MK181220**
IPSC: 377.4 pA
Latency: 0.93 ms
Failure rate: 0%

8 contacts

Interneuron

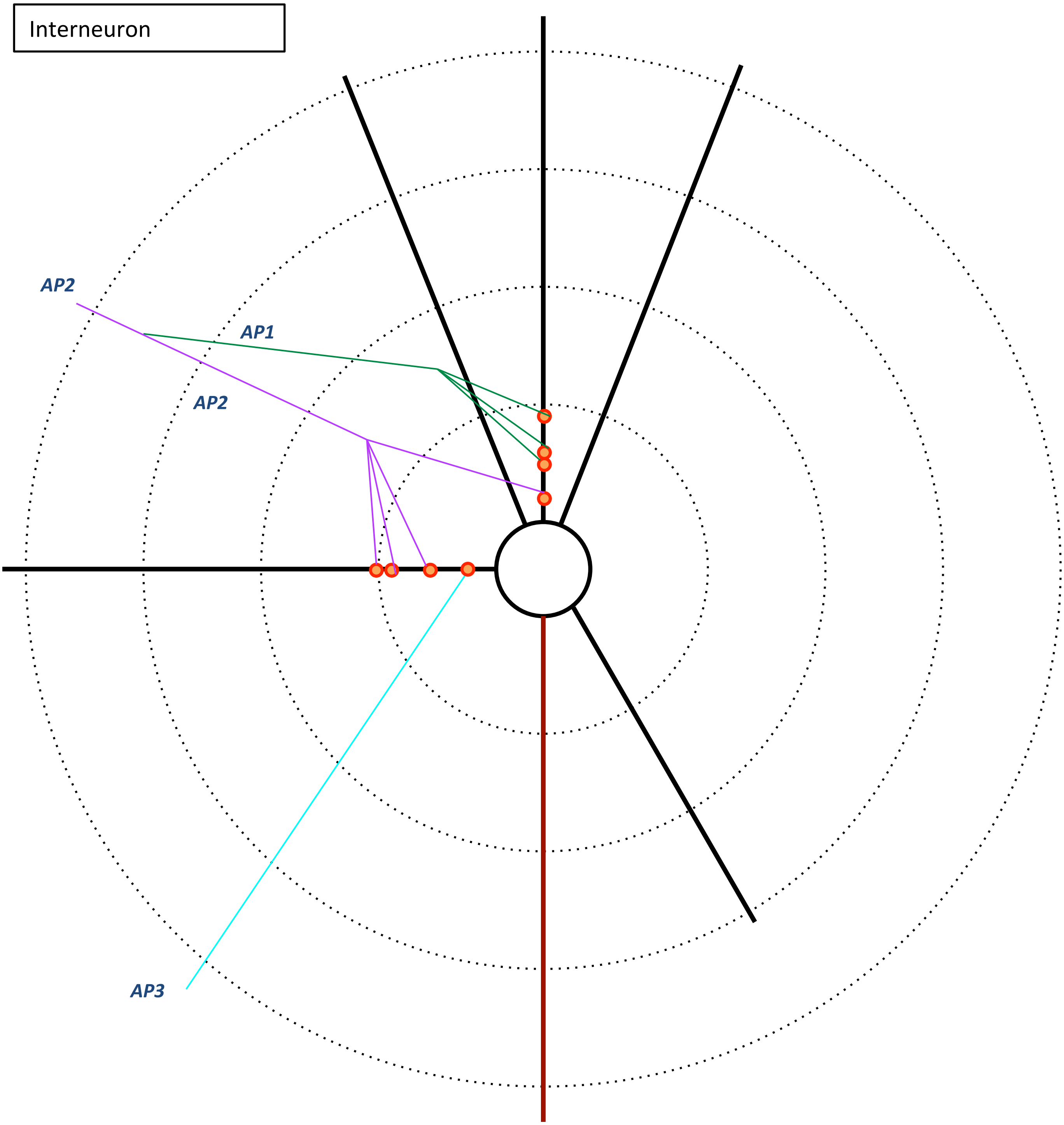

**MK181127**
IPSC: 61.8 pA
Latency: 0.87 ms
Failure rate: 0%

8 contacts

Interneuron

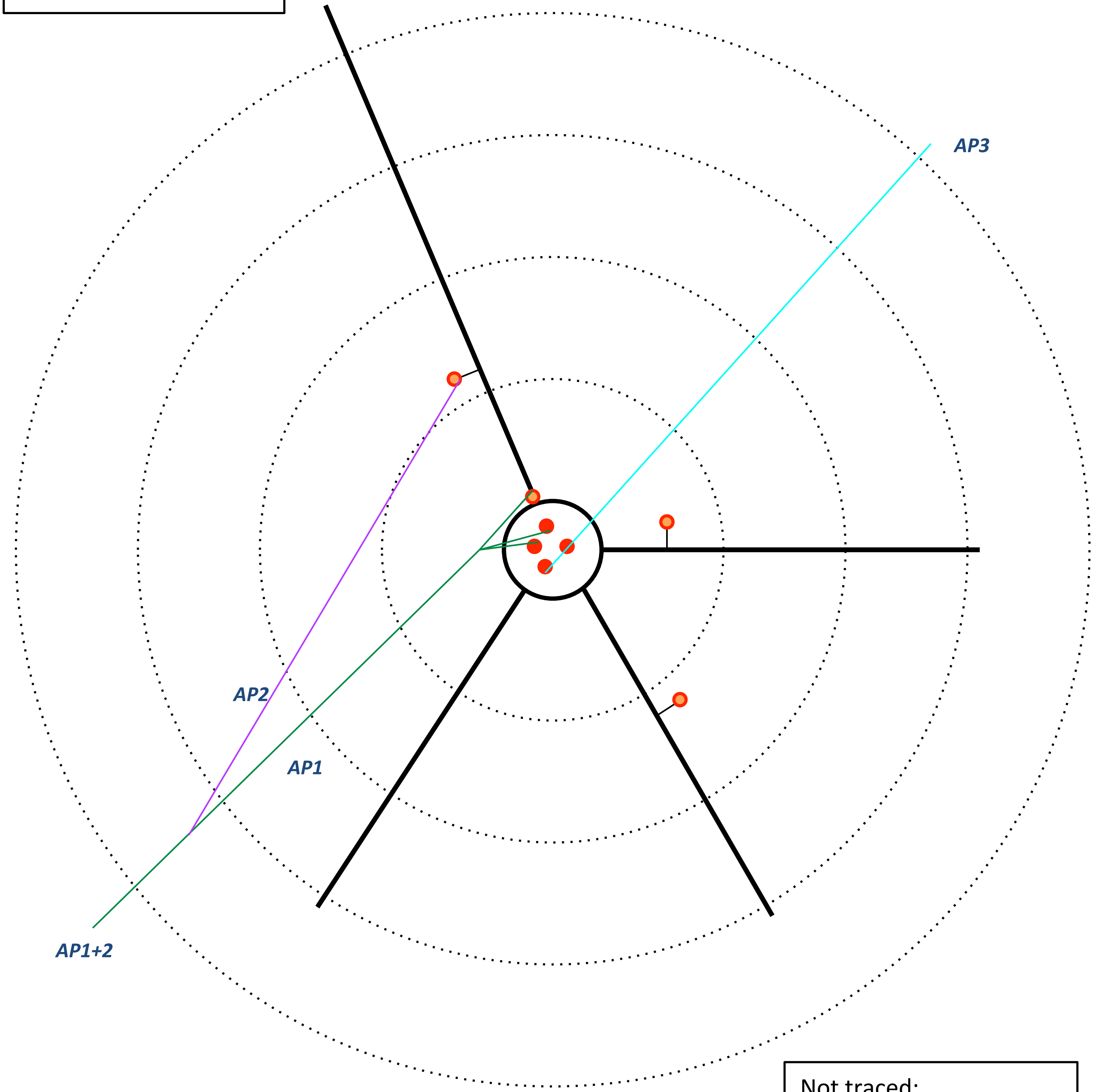


Not traced:
2 more contacts on proximal dendrites

**MK181221-1**
IPSC: 417.1 pA
Latency: 1.32 ms
Failure rate: 0 %

15 contacts

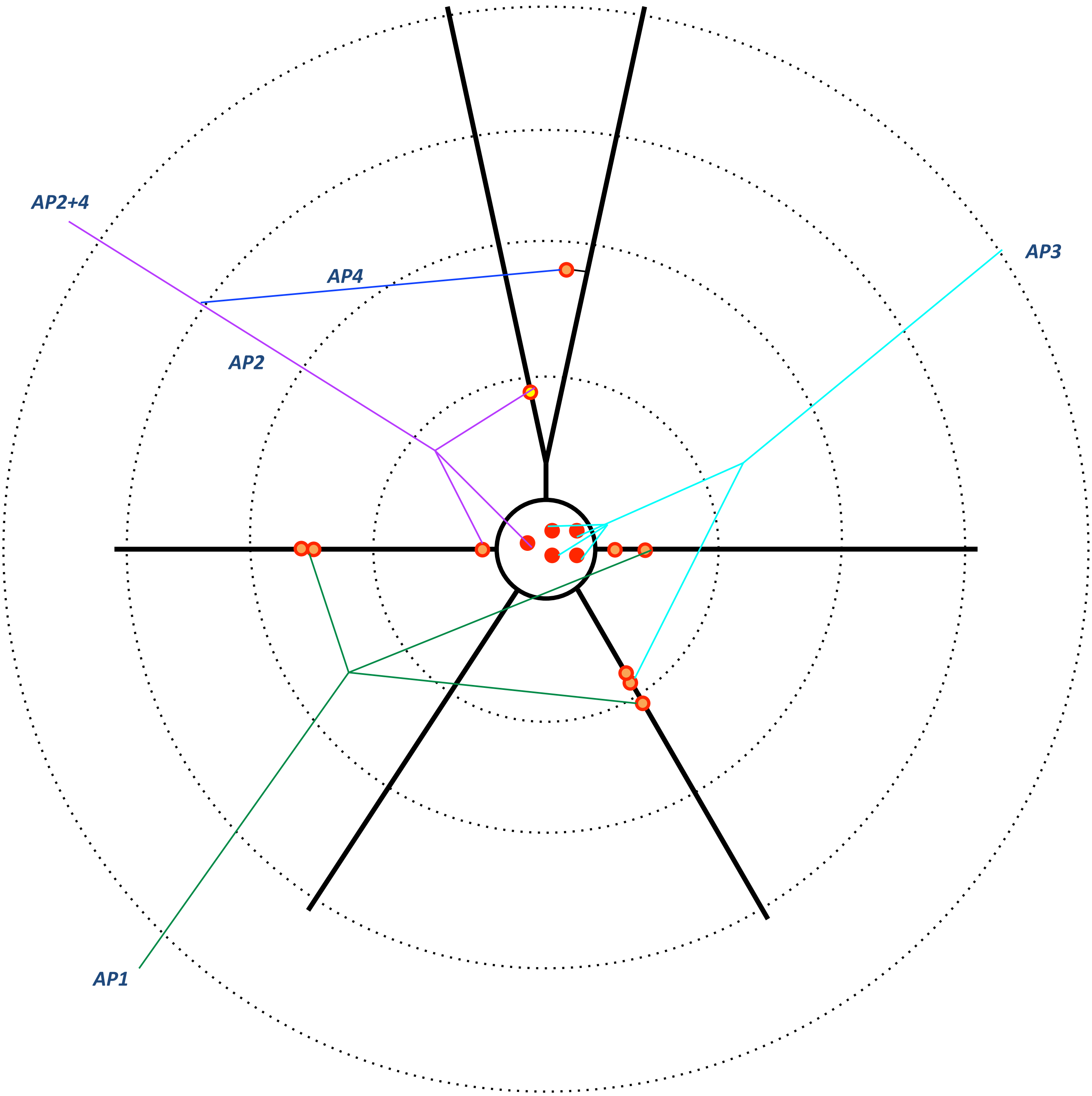

**MK190109-2**
IPSC: 21.1 pA
Latency: 1.64 ms
Failure rate: 48.48%

8 contacts

No change
Ampl: 21.06
Lat: 1.64

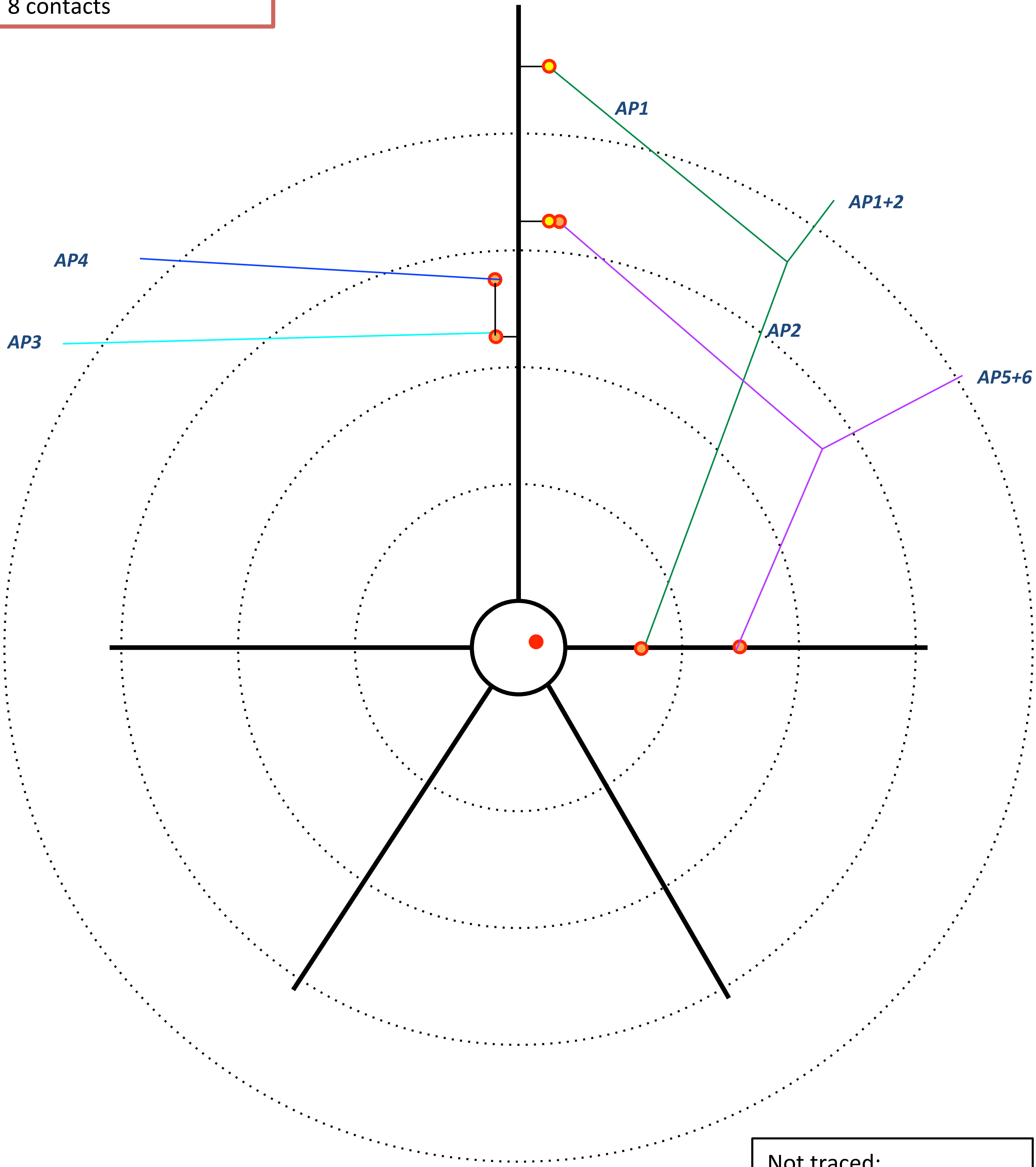


Not traced:
1 contact on cell body

**MK180817**
IPSC: 223.4 pA
Latency: 2.46 ms
Failure rate: 0%

14 contacts

**Axonal paths not traced**

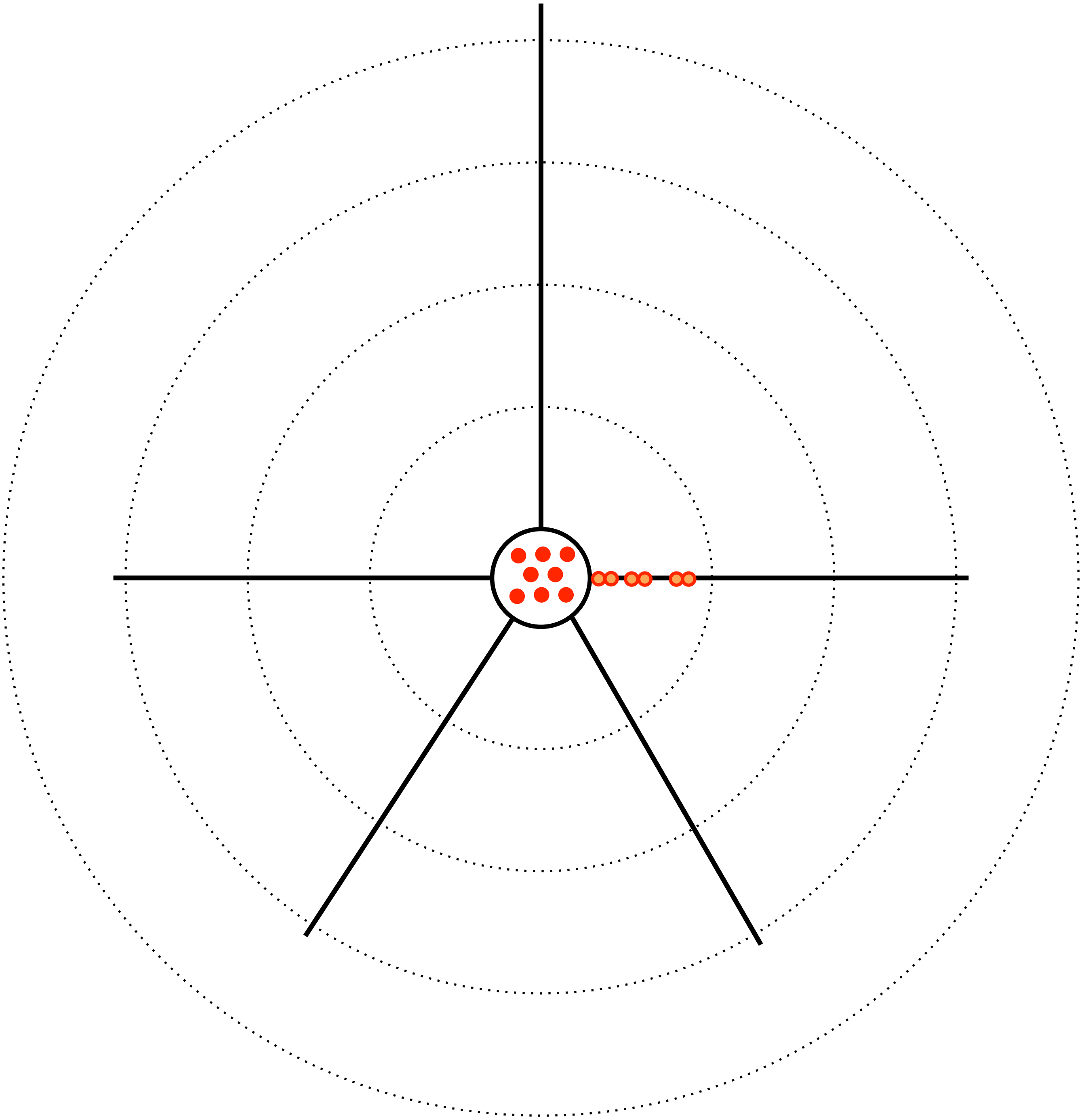
